# Structural basis of transcription-translation coupling and collision in bacteria

**DOI:** 10.1101/2020.03.01.971028

**Authors:** Michael William Webster, Maria Takacs, Chengjin Zhu, Vita Vidmar, Ayesha Eduljee, Mo’men Abdelkareem, Albert Weixlbaumer

## Abstract

Prokaryotic messenger RNAs (mRNAs) are translated as they are transcribed. The pioneering ribosome potentially contacts RNA polymerase (RNAP), forming a supramolecular complex known as the expressome. The basis of expressome assembly and its consequences for transcription and translation are poorly understood. Here we present a series of structures representing uncoupled, coupled and collided expressome states determined by electron cryomicroscopy. A bridge between the ribosome and RNAP can be formed by the transcription factor NusG, stabilizing an otherwise variable interaction interface. Shortening of the intervening mRNA causes a substantial rearrangement that aligns the ribosome entrance-channel to the RNAP exit-channel. In this collided complex, NusG-linkage is no longer possible.

These structures reveal mechanisms of coordination between transcription and translation and provide a framework for future study.

**One Sentence Summary:** Structures of the molecular assembly executing gene expression shed light on transcription translation coupling.

## Introduction

All organisms express genetic information in two steps. Messenger RNAs (mRNAs) are transcribed from DNA by RNA polymerase (RNAP), and then translated by ribosomes to proteins. In prokaryotes, translation begins as the mRNA is synthesized, and the pioneering ribosome on an mRNA is spatially close to RNAP (*1, 2*). Coordination of transcription with translation regulates gene expression and prevents premature transcription termination (*3, 4*). The trailing ribosome inhibits RNAP backtracking, contributing to synchronization of transcription and translation rates *in vivo* (*5, 6*).

Coordination may also involve physical contacts between RNAP and the ribosome. The conserved transcription factor NusG binds RNAP through its N-terminal domain (NusG-NTD), and ribosomal protein uS10 through its C-terminal domain (NusG-CTD) both *in vitro* and *in vivo* (*7, 8*). While a bridge may be formed by simultaneous binding, the consequences of this are unknown. RNAP and the ribosome also interact directly (*9-11*). A transcribing-translating ‘expressome’ complex formed by collision of ribosomes with stalled RNAP in an *in vitro* translation reaction was reconstructed at 7.6 Å resolution (*9*). Yet this architecture does not permit a NusG-mediated bridge.

### Uncoupled Expressome

We sought to structurally characterize mechanisms of transcription-translation coupling, and resolve the relationship between NusG and the collided expressome. Expressomes were assembled by sequential addition of purified *Escherichia coli* (*E. coli*) components (70S ribosomes, tRNAs, RNAP and NusG) to a synthetic DNA-mRNA scaffold (Fig S1A-C). An mRNA with 38 nucleotides separating the RNAP active site from the ribosomal P-site was chosen to imitate a state expected to precede collision (*12*).

A reconstruction of the expressome was obtained at 3.0 Å nominal resolution by electron cryomicroscopy (cryo-EM) (Fig 1A and S1D,E). Yet RNAP and ribosome do not adopt a single relative orientation within the expressome, and focused refinement was required to attain a reconstruction of the RNAP region at 3.8 Å nominal resolution (Fig 1A and S2). Refined atomic models collectively present the key steps of prokaryotic gene expression within a single molecular assembly (Fig 1B).

**Fig. 1.**
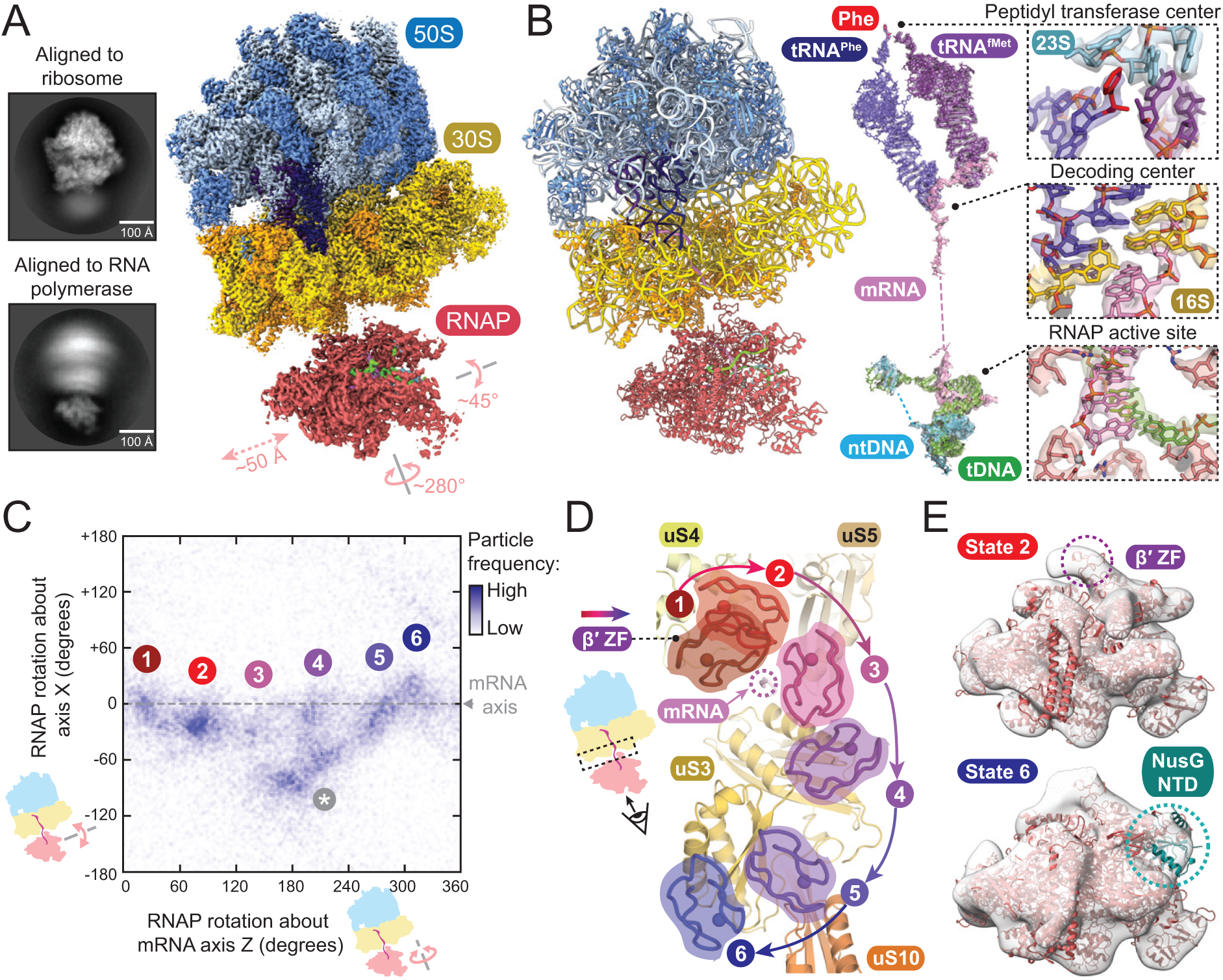
Structural models of the uncoupled expressome. **(A)** Representative cryo-EM 2D class averages showing conformational variability (left), and cryo-EM maps of ribosome and RNAP in the uncoupled expressome (right). RNAP is shown in position 2 (see panel E), with measured rotation and translations of RNAP indicated. **(B)** Atomic model of the uncoupled expressome in ribbon representation (left), and the central steps in gene expression shown by segmented cryo-EM maps with superimposed atomic coordinates (right). **(C)** Plot of RNAP-70S relative orientation with clusters indicating a series of states (1-6) distinguished by rotation of RNAP. One state (*) is likely not populated in a physiological context of longer DNA (see Fig. S3B). **(D)** Position of the RNAP β′-ZF in each expressome model relative to the ribosome surface. **(E)** NusG is present in state 6 but not in state 2. Focused cryo-EM maps shown filtered to 20 Å resolution with fitted coordinates.

Direct contacts between RNAP and the ribosome, if they occur, are not stable in this complex and the mRNA is the only consistent connection. We characterized the dynamics of the complex by plotting the range of RNAP positions relative to the ribosome using the angular assignments of particles from focused reconstructions (Fig 1C and Fig S3A). RNAP is loosely restrained to a plane perpendicular to an axis connecting the RNAP mRNA exit-channel to the ribosomal mRNA entrance-channel (Movie S1). Within this plane, RNAP rotates freely, with clusters representing a series of preferred relative orientations (Fig 1C and Fig S3A,B).

RNAP and ribosome models could be placed in reconstructions generated from particles in each cluster (Fig S3C-F). These expressome models collectively suggest a continuous movement of RNAP along the ribosome surface involving substantial changes in both rotation (∼280°) and translation (∼50 Å) (Fig 1D and Movie S1). The closest domain of RNAP to the ribosome is the zinc finger of the β′ subunit (β′-ZF) in all models. In states 1-3, β′-ZF sits within a funnel-shaped depression between the head, body, and shoulder domains of the 30S subunit bounded by ribosomal proteins uS3, uS4 and uS5. We estimate RNAP transits from state 1 through states 2-5 to reach the position shown by model 6. Here, the RNAP β′-ZF is between uS3 and uS10 on the 30S head domain.

NusG-NTD is bound to RNAP in expressome state 6, but not states 1 and 2 (Fig 1E). We determined that a substantial fraction of the imaged particles were lacking NusG due to dissociation during gradient purification (Fig S3G). Importantly, the predicted position of the NusG-CTD bound to uS10 is close to the NusG-NTD bound to RNAP only in state 6.

### Coupled Expressome

An improved reconstruction of the NusG-coupled expressome was obtained from a new sample prepared with increased NusG occupancy (Fig 2A and Fig S4A, B). Heterogeneity in the position of ribosome and RNAP was substantially reduced, but focused refinement was required to obtain well-resolved ribosome and RNAP reconstructions (3.4 Å and 7.6 Å respectively) (S4C-E). Continuous density in the unfocused reconstruction confirmed NusG bridges RNAP and the ribosome (Fig. 2A). We constructed an atomic model of the NusG-coupled expressome by fitting and refining a ribosome model, and docking a published RNAP-NusG-NTD model consistent with our map (*13*) into their consensus positions in the unfocused reconstruction (Fig 2B).

**Fig. 2.**
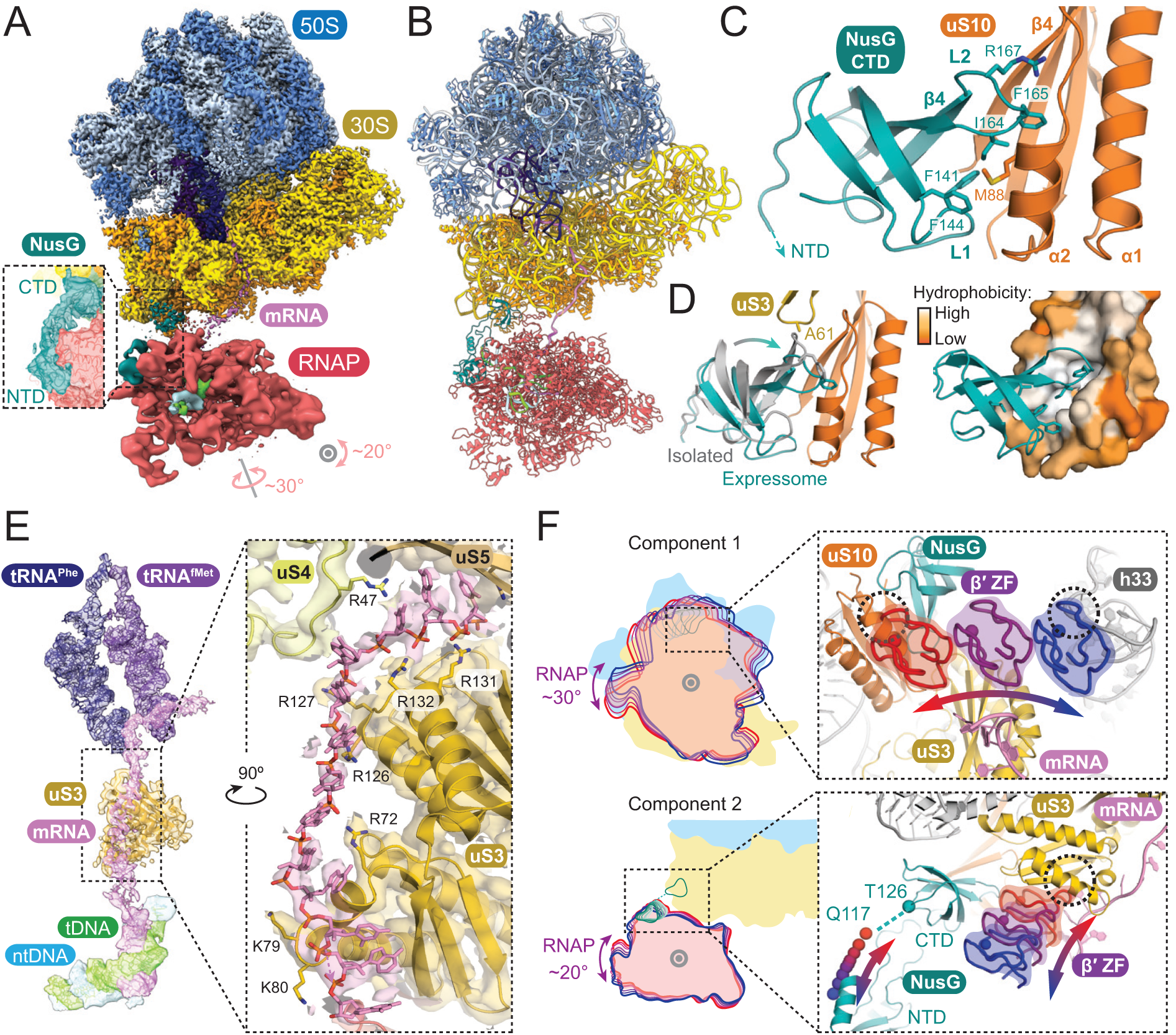
Structural models of the NusG-coupled expressome. **(A)** Cryo-EM maps of ribosome and RNAP in the coupled expressome. Inset shows continuous electron density between the NusG NTD and CTD domains in unfocused map filtered to 8 Å (slice view). **(B)** Ribbon representations of the NusG-coupled expressome model. **(C)** Interaction of NusG-CTD with ribosomal protein uS10. **(D)** Structural superposition with the isolated NusG-uS10 complex based on alignment to uS10 (grey; PDB code 2KVQ) (left), and hydrophobic pocket created by conformational change of uS10 (right). **(E)** mRNA connecting the ribosome mRNA entrance-channel to RNAP exit-channel shown by cryo-EM map filtered to 4 Å and fitted model. **(F)** The range of RNAP positions relative to the ribosome surface determined by multi-body refinement. Cartoon of two principal components accounting for 44% of variance (left). Component 1 involves rotation in a plane approximately parallel to the surface of the ribosome and is limited by clashes between the β′-ZF of RNAP and either uS10 or h33 (dashed circles). Component 2 is an orthogonal rotation limited by extension of the flexible NusG linker (residues Q117-T126) in one direction and clash between β′-ZF and uS3 in the other (dashed circle). Positions of RNAP β′-ZF and NusG residue Q117 indicate trajectories (red through purple to blue).

Additional density corresponding to the NusG-CTD bound to uS10 was identified on the ribosome, which otherwise closely resembled that of the uncoupled expressome. The NusG-CTD is a KOW domain, consisting of a five-stranded β-barrel. As in the isolated NusG-uS10 complex determined by NMR (*7*), strand β4 of NusG aligns with strand β4 of uS10, forming an extended intermolecular β-sheet (Fig 2C). Yet NusG and uS10 are significantly closer in the expressome than in isolation because NusG loops L1 (F141 and F144) and L2 (I164, F165, R167) insert into a hydrophobic pocket of uS10 that is enlarged by movement of helix α2 (Fig 2D and S5A-D). F165 of NusG in particular is embedded within uS10. This accounts for its crucial role in binding uS10 identified by mutational studies (*8*). The altered position of NusG not only increases the area contacting uS10 but avoids clashing with neighboring ribosomal protein uS3 (Fig 2D).

The NusG-CTD recruits Rho to terminate synthesis of untranslated mRNAs (*14*). In the coupled expressome, NusG binds uS10 with the same interface it binds Rho, suggesting the events are mutually exclusive (Fig S5E) (*15*). The structure of the expressome thereby explains how the trailing ribosome is sensed by NusG, and transcription termination is consequently reduced.

Binding of the NusG-NTD to RNAP suppresses backtracking by stabilizing the upstream DNA duplex (*13, 16*). In the expressome, space for the upstream DNA is further restricted by an extended channel formed by uS10 and NusG. The interaction of the NusG-CTD with uS10 is predicted to reduce dissociation of NusG-NTD from RNAP through increased avidity (*17*). The RNAP-NusG complex within the coupled expressome is likely stabilized by the trailing ribosome, and transcription elongation is consequently favored.

The mRNA exit-channel of RNAP is separated from the entrance-channel of the ribosome by ∼60 Å. Continuous electron density on the solvent side of uS3 allowed modeling of the intervening 12 mRNA nucleotides, completing the mRNA path from synthesis to decoding (Fig 2E and Fig S6A-C). The interpretability of the electron density varies considerably, however, and this model is considered one of an ensemble of mRNA conformations.

The RNAP mRNA exit-channel is adjacent to uS3 residues R72, K79 and K80, and clear electron density for mRNA in this region suggests a relatively stable contact. The path continues to four arginines immediately outside the ribosomal mRNA entrance-channel (R126, R127, R131 and R132). R131 and R132 were previously identified as imparting ribosomal helicase activity (*18*). The mRNA path in this region is close to, but different from, that observed previously in structures of mRNA-bound ribosomes (*19*) (Fig S6D-F).

Binding of the nascent transcript by uS3 likely modulates secondary structure formation. Structured mRNAs can decrease translation rates (*20*), stabilize transcriptional pauses (e.g. the *E. coli his*-pause (*21*)) or induce transcription termination (*22*). While the ribosome can unwind mRNA secondary structure with basic residues in the mRNA entrance-channel (*18*), preventing mRNAs folding downstream likely aids translation efficiency. We propose that by positioning RNAP in line with an extended series of basic residues, NusG supports nascent mRNAs staying single-stranded and thereby enhances both transcription and translation efficiency.

No stable contacts are observed between the core subunits of RNAP and the ribosome in the NusG-coupled expressome. The relative position of RNAP and the ribosome varies between particles, albeit substantially less than the sample with partial NusG occupancy (Fig S4A,B). Analysis of movement by multi-body refinement (*23*) revealed RNAP is constrained by the tethering action of NusG and the insertion of β′-ZF into a cavity formed by uS3, uS10, NusG and helix 33 of 16S rRNA (h33) (Fig 2F and Movie S2).

### Collided Expressome

The mRNA spanning the mRNA exit- and entrance-channels is in an extended conformation in the NusG-coupled expressome. To test if coupling by NusG is possible when the spanning mRNA is shorter, we obtained a reconstruction of a NusG-containing expressome with an mRNA shortened to 34 nucleotides between the ribosomal P-site and the RNAP active site (Fig 3A and S7). A model was constructed as described for the coupled expressome (Fig 3B).

**Fig. 3.**
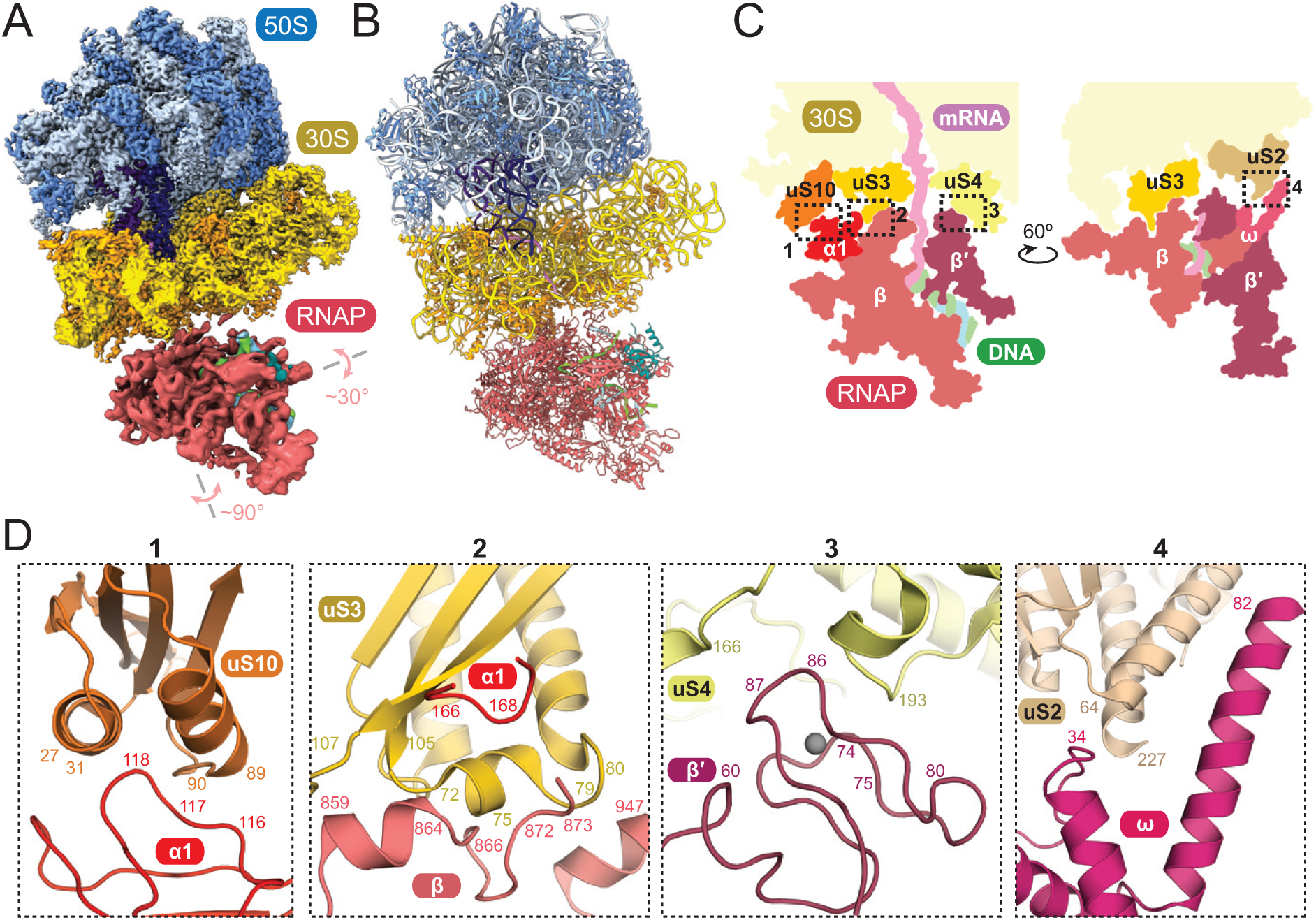
Structural models of the collided expressome. **(A-B)** Cryo-EM maps and model of the collided expressome. **(C)** Schematic cross-section indicating four regions of close contact between RNAP and ribosome. **(D)** Details of the interaction interfaces of RNAP with ribosome.

RNAP is positioned close to the ribosome mRNA entrance-channel, more than 50 Å from its location in the NusG-coupled expressome. Consistent with this change in position, RNAP still binds the NusG-NTD but is no longer tethered through the NusG-CTD to uS10. We determined the structure of an equivalent sample lacking NusG, and confirmed the position of RNAP is very similar (Fig S8A). The architecture is therefore not NusG-dependent, and is similar to that of particles from clusters 1 and 2 of the uncoupled expressome (Fig S8B). We conclude that coupling of NusG to the ribosome requires the P-site to be more than 34 nucleotides from the 3’ end of the mRNA.

The arrangement of RNAP and ribosome in our structure resembles the expressome formed by collision of translating ribosomes with stalled RNAP (RNAP backbone RMSD ∼3Å based on 16S rRNA superposition) (*9*) (Fig S8B and S9A,B). We therefore term this molecular state the ‘collided expressome’. The reconstruction previously reported was resolved to 7.6 Å, and our improved model allowed us to define the interaction surfaces of RNAP and the ribosome in more detail.

Four regions are in close proximity: uS10 with the N-terminal domain of the RNAP α1 subunit, uS3 with RNAP subunits α1 and the β-flap domain, uS4 with β′-ZF, and uS2 with RNAP ω-subunit (Fig 3C,D and S8C-D). The contacts bury a total surface area of ∼3000 Å^2^. Yet RNAP moves relative to the ribosome, albeit less than in the samples previously analyzed (Fig S8B). The contacts between RNAP and ribosome are likely transient, and the contact area consequently variable. The observed RNAP-ribosome configuration allows striking structural complementarity between the molecular surfaces.

Rotation of RNAP relative to the ribosome beyond the observed position would cause steric clashes (Fig 4A and S10). We hypothesized that the architecture of the collided expressome is the product of structural complementarity and the energetically-favorable minimization of mRNA path length. To test this, we generated ∼18,000 hypothetical expressome models representing an exhaustive search of RNAP rotations about the mRNA axis at a series of distances along it (2° rotational step size, 0.5 Å translational step size). After excluding clashing models, we found that the shortest mRNA path is achieved by the RNAP orientations observed by cryo-EM (Fig 4B). A simple model is therefore sufficient to explain the observed orientation of RNAP relative to the ribosome: when inserting into the mRNA entrance-channel cavity on the ribosome, RNAP adopts orientations with the greatest structural complementarity to allow the intervening mRNA to span the shortest distances.

**Figure 4:**
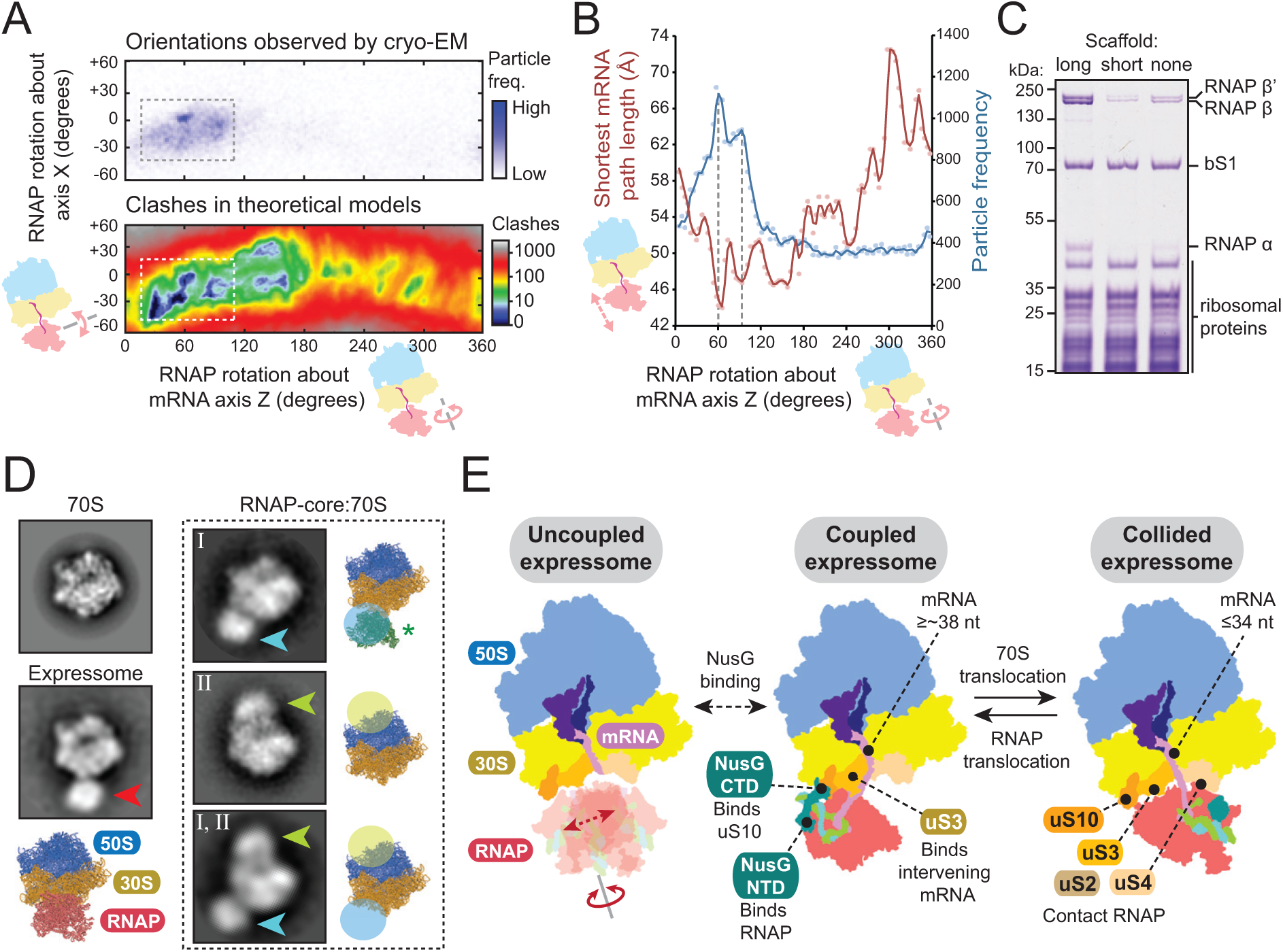
Formation of RNAP-70S complexes. **(A)** Collided expressome RNAP-70S relative orientations observed by cryo-EM (top) correspond to a restricted space that avoids steric clashes (bottom). **(B)** The most common RNAP positions in the collided expressome (blue line) coincide with minima of the intervening mRNA path length (red line). **(C)** Gradient co-purification of RNAP with 70S ribosomes depends on the nucleic acid scaffold. RNAP-70S complexes were formed under low-salt conditions using an mRNA long enough to allow ribosome binding (‘long’), or not (‘short’), or no nucleic acids (‘none’). Coomassie-stained SDS-PAGE of ribosome-containing sucrose gradient peak shown. **(D)** Negative stain EM class averages of 70S-RNAP complexes show distinct binding sites for core RNAP sample (cyan and lime arrows) compared with expressome sample (red arrow). Position of RNAP from 30S-RNAP complex superimposed (green asterisk). **(E)** Key features and interchange between expressome complexes during transcription-translation coordination. In the uncoupled expressome, RNAP is loosely restrained and adopts various orientations. Coupling by NusG aligns the mRNA with ribosomal protein uS3 and restricts the position of RNAP. Once the ribosome approaches RNAP further, the collided state forms in which the mRNA length is limiting and NusG no longer links the two machineries.

### Alternative RNAP ribosome interactions

We sought to clarify whether expressome formation is driven by concurrent binding to the same mRNA, or if specific ribosome-RNAP contacts contribute. Co-purification of RNAP with ribosomes was substantially reduced when the mRNA did not support concurrent ribosome binding. Yet RNAP lacking DNA or mRNA (‘RNAP-core’) bound ribosomes more stably (Fig 4C and S9C,D). This has been observed previously, and it was thought expressome formation can be mRNA-independent (*9, 11*).

To examine this, we imaged samples assembled without further purification and lacking nucleic acid scaffolds (core RNAP-70S) by negative stain EM. While no expressomes formed, confirming the importance of an mRNA that supports binding of both machineries, we observed at least two alternative RNAP binding sites (Fig 4D). The sites can be described only approximately from this data, but one (site I) is consistent with an interaction with ribosomal protein uS2 observed in a core RNAP-30S complex (*10*). Saturation of ribosomes with ribosomal protein bS1, which has no effect on expressome formation (Fig S11A), abolished occupancy of site I without affecting the second site (site II). Addition of a nucleic acid scaffold containing just a short mRNA (minimal scaffold) abolished occupancy of site II only, while addition of both (short mRNA scaffold, bS1) abolished both (Fig S11). Further analysis is required to assess the potential biological role, but the existence of additional 70S-RNAP interaction modes highlights the complexity of possible transcription-translation connections.

## Conclusions

Thus, the expressome is mRNA-linked and consequently dynamic. A level of structural independence may be required for the internal movements that occur during the reaction cycle of each complex. Coupling by NusG restrains RNAP motions, and likely also happens when the two machineries are more separated than in the complexes described here, but not when they collide (Fig 4E). Importantly, translation factor binding is compatible with all the observed RNAP orientations. This study provides a basis for future work on the role of coupling in gene expression, and its regulation by transcription factors and regulatory mRNA structures.

## Materials and Methods

### Materials

For plasmid construction, we used the *E. coli* TOP10 strain (Invitrogen). For recombinant protein expression, we constructed an *E. coli* strain, called LACR II (<u>L</u>ow <u>A</u>bundance of <u>C</u>ellular <u>R</u>Nases). LACR II is derived from the *E. coli* LOBSTR strain (*24*) with additional RNase deletions to lower the amount of RNAse contamination in purified protein samples (details to be published elsewhere).

*E. coli* strain BL21(DE3)rpoA-HRV3C-CTD-(His)_10_ (*25*) with a human rhinovirus 3C (HRV3C) protease cleavage site in the linker between the α subunit C-terminal (CTD), and N-terminal domains (NTD) was a generous gift from the Darst lab.

*E. coli* strain HMS174 and plasmids to express *E. coli* tRNA^fMet^ and *E. coli* tRNA^Phe^ were generous gifts from the Ramakrishnan lab.

Plasmid pAX1_(His)_10_-TwinStrep-HRV3C-rpsA for expression of *E. coli* small ribosomal subunit protein bS1 was constructed by amplification of the *E. coli rpsA* gene with primers 5′-GTTCTGTTTCAGGGTCCGCATATGACTGAATCTTTTGCTCAACTCTTTG-3′ and 5′-AGTGGTGGTGGTGGTGGTGCTCGAGTTACTCGCCTTTAGCTGCTTTGAAAGC-3′ and insertion into pAX1_(His)_10__TwinStrep_HRV3C at the NdeI and XhoI sites using the SLiCE method {Zhang:2012bn}.

Plasmid pAX0_(His)_10_-HRV3C-pheS_pheT for expression of *E. coli* Phenylalanyl-tRNA synthetase (PheRS) was constructed by amplification of the *E. coli pheS* and *pheT* genes with primers 5′-GTTCTGTTTCAGGGTCCGCATATGTCACATCTCGCAGAACTGGTTGC-3′ and 5′-AGTGGTGGTGGTGGTGGTGCTCGAGTCAATCCCTCAATGATGCCTGGAATCG-3′ and insertion into pAX0_(His)_10__HRV3C at the NdeI and XhoI sites using the SLiCE method.

### E. coli RNAP

*E. coli* RNA polymerase (RNAP) with a C-terminally (His)_10_-tagged β′-subunit was overexpressed in *E. coli* LACR II strain from pVS11_rpoA_rpoB_rpoC-HRV3C-(His)_10__rpoZ co-transformed with pACYC_Duet1_rpoZ to avoid substoichiometric amounts of the RNAP ω-subunit. *E. coli* RNAP with cleavable, (His)_10_-tagged α-subunit C-terminal domains (*E. coli* RNAP_Δα-CTD) was overexpressed in BL21(DE3)rpoA-HRV3C-CTD(His)_10_ from pVS10_rpoA-HRV3C-CTD-(His)_10__rpoB_rpoC_rpoZ co-transformed with pACYC_Duet1_rpoZ. Both RNAPs were purified as described before (*26*). For expression, 12 L of LB culture (100 µg/ml Ampicillin, 34 µg/ml Chloramphenicol) were induced at an OD_600_ of 0.6-0.8 with 0.5 mM IPTG overnight at 18°C. Cells were harvested by centrifugation, resuspended in 5 volumes of lysis buffer (50 mM Tris-HCl, pH 8.0, 5% glycerol, 1 mM EDTA, 10 µM ZnCl_2_, 10 mM DTT, 0.1 mM PMSF, 1 mM benzamidine, DNase I (0.1 mg/50g cell), EDTA-free protease inhibitor cocktail (Sigma-Aldrich cOmplete, 1 tablet/50 ml)) and lysed by sonication. The lysate was cleared by centrifugation at 40,000 g for 30 minutes. RNAP was isolated from the supernatant by polyethyleneimine fractionation followed by ammonium sulfate precipitation as described previously (*27*). The precipitate was resuspended in IMAC binding buffer (20 mM Tris-HCl, pH 8.0, 1 M NaCl, 5% glycerol, 10 µM ZnCl_2_, 5 mM β-mercaptoethanol, 0.1 mM PMSF, 1 mM benzamidine), loaded on a 20 ml Ni-IMAC Sepharose HP column (GE Healthcare) and after several washing steps RNAP was eluted into IMAC elution buffer (IMAC binding buffer containing 250 mM imidazole). Peak fractions were pooled, and dialyzed overnight in the presence of His-tagged HRV3C (PreScission) protease (1 mg HRV3C per 8 mg of protein) into dialysis buffer (20 mM Tris-HCl, pH 8.0, 1 M NaCl, 5% glycerol, 5 mM β-mercaptoethanol, 10 µM ZnCl_2_). Cleaved RNAP was separated from uncleaved RNAP and HRV3C protease by subtractive Ni-IMAC. The sample was then dialyzed into TGE buffer supplemented with ZnCl_2_ (10 mM Tris-HCl, pH 8.0, 5% glycerol, 0.1 mM EDTA, 10 µM ZnCl_2_, 1 mM DTT, 0.1 mM PMSF, 1 mM benzamidine) until the conductivity was ≤ 10 mS/cm. RNAP was then loaded on a 50 ml Bio-Rex 70 column (Bio-Rad) equilibrated with Bio-Rex buffer (10 mM Tris-HCl, pH 8.0, 5% glycerol, 0.1 mM EDTA, 0.1 M NaCl, 10 µM ZnCl_2_, 1 mM DTT, 0.1 mM PMSF, 1 mM benzamidine) and was eluted using a linear gradient over 5 column volumes into Bio-Rex buffer containing 1 M NaCl. The peak was concentrated and further purified by size-exclusion chromatography using a HiLoad Superdex 200 PG 26/600 column (GE Healthcare) equilibrated with GF buffer (10 mM HEPES, pH 8.0, 0.5 M KCl, 1% glycerol, 10 µM ZnCl_2_, 1 mM MgCl_2_, 2 mM DTT, 0.1 mM PMSF, 1 mM benzamidine). The final protein was dialyzed into EM buffer (10 mM HEPES, pH 8.0, 150 mM KOAc, 5 mM Mg(OAc)_2_, 10 µM ZnCl_2_, 2 mM DTT), concentrated to approximately 80 mg/ml and aliquots were flash frozen and stored at −80°C.

### E. coli NusG

*E. coli* NusG with an N-terminal (His)_6_-tag was overexpressed in *E. coli* LACR II strain from pSKB2_(His)_6_-HRV3C-NusG. For expression, 6 L of LB culture (50 µg/ml Kanamycin) was induced at an OD_600_ of 0.6 with 1 mM IPTG for 3 hours at 37°C. Cells were harvested by centrifugation, resuspended in 4 volumes of lysis buffer (50 mM Tris-HCl pH 8.0, 2 mM EDTA, 233 mM NaCl, 5% glycerol, 5 mM β-mercaptoethanol, 0.1 mM PMSF, 1 mM benzamidine, EDTA-free protease inhibitor cocktail (Sigma-Aldrich cOmplete, 1 tablet/50 ml)) and lysed by sonication. The lysate was cleared by centrifugation at 40,000 g for 30 minutes at 4°C. The nucleic acids and their interacting proteins were precipitated by adding 0.6% of polyethyleneimine and removed by centrifugation at 45,000 g for 20 minutes at 4°C. (NH_4_)_2_SO_4_ was added to the supernatant to a final concentration of 0.37g/ml and the precipitate was collected by centrifugation at 45,000 g for 20 minutes at 4°C. The pellet was resuspended in IMAC buffer A (50 mM Tris-HCl, pH 8.0, 0.5 M NaCl, 5 mM imidazole, 1 mM β-mercaptoethanol, 0.1 mM PMSF, 1 mM benzamidine) and loaded on a 5 ml HiTrap IMAC HP column (GE Healthcare). After a wash step for 3 column volumes in 30% IMAC buffer B (same as IMAC A but 500 mM imidazole), NusG was eluted at 40% of IMAC buffer B (200 mM imidazole). Peak fractions were pooled, and dialyzed overnight in the presence of His-tagged HRV3C (PreScission) protease (1 mg HRV3C per 18 mg of protein) into dialysis buffer (50 mM Tris-HCl, pH 8.0, 0.25 M NaCl, 5% glycerol, 1 mM β-mercaptoethanol). Cleaved NusG was separated from uncleaved NusG and HRV3C protease by subtractive Ni-IMAC. The sample was then dialyzed into ion-exchange buffer A (10 mM Tris-HCl, pH 8.0, 0.1 mM EDTA, 5% glycerol, 1 mM DTT, 0.1 mM PMSF, 1 mM benzamidine). NusG was loaded on a 5 ml HiTrap Q HP column (GE Healthcare) and eluted using a gradient of 0-100% ion-exchange buffer B (10 mM Tris-HCl, pH 8.0, 0.2 M NaCl, 0.1 mM EDTA, 5% glycerol, 1 mM DTT, 0.1 mM PMSF, 1 mM benzamidine) for 20 column volumes. The peak was concentrated and further purified by size-exclusion chromatography using a Superdex 75 16/600 column equilibrated with GF buffer (10 mM Tris-HCl, pH 8.0, 0.5 M NaCl, 0.1 mM EDTA, 5% glycerol, 1 mM DTT, 0.1 mM PMSF, 1 mM benzamidine). The final protein was concentrated to 5 mg/ml and aliquots were flash frozen and stored at −80°C.

### E. coli 70S ribosome

Tight-coupled 70S ribosomes were purified from *E. coli* strain LACR II following standard procedures (*28, 29*). S1-depleted 70S ribosomes were prepared using immobilized poly-U chromatography (*30*). 70S ribosomes were dissociated into 30S and 50S subunits and the purified subunits were then reassociated to form S1-free 70S ribosomes (*29*). The complete purification of *E. coli* 70S ribosomes was done at 0-4°C and all buffers contained 6 mM β-mercaptoethanol, 1 mM benzamidine and 0.1 mM PMSF added just before use. Briefly, *E. coli* LACR II cells were grown at 37°C in LB until they reached an OD_600_ of 1.3. The harvested cells were resuspended in buffer A (20 mM Tris-HCl, pH 7.5, 10.5 mM Mg(OAc)_2_, 100 mM NH_4_Cl, 0.5 mM EDTA, DNase I (0.4 mg/50g cell), protease inhibitor cocktail (Sigma-Aldrich cOmplete, 1 tablet/50 ml); 3-5 ml/g cell paste), lysed by sonication and the cell lysate was centrifuged in a Beckman Type 45 Ti rotor for 1 hour at 185,000 g. After the centrifugation, the clear top part of the supernatant was carefully taken, filtered through a 0.22 µm membrane and layered on 25 ml sucrose cushion (20 mM Tris-HCl, pH 7.5, 1.1 M sucrose, 1 M NH_4_Cl, 10.5 mM Mg(OAc)_2_, 0.5 mM EDTA) in 45 Ti tubes (40 ml supernatant on 25 ml cushion/tube). The ribosomes were sedimented overnight at 185,000 g for 18 hours. The pellet was washed and resuspended in buffer C (20 mM Tris-Cl, pH 7.5, 1 M NH_4_Cl, 10.5 mM Mg(OAc)_2_, 0.5 mM EDTA) and sedimented through an additional sucrose cushion. To isolate tightly coupled 70S ribosomes and to remove excess 50S and 30S subunits the pellet was washed and resuspended in buffer D (20 mM Tris-HCl, pH 7.5, 60 mM NH_4_Cl, 6 mM Mg(OAc)_2_, 0.25 mM EDTA) and loaded on 15–30% sucrose gradient. This gradient was centrifuged in an SW28 rotor at 58,000 g for 18 hours. The gradient was fractionated and the peak containing 70S ribosomes were collected avoiding any contamination by 50S subunits. The pooled fractions were diluted 2-fold with buffer E (20 mM Tris-HCl, pH 7.5, 60 mM NH_4_Cl, 20 mM Mg(OAc)_2_, 0.25 mM EDTA) and were again sedimented overnight at 185,000 g for 18 hours in a Beckman Type 45 Ti rotor. The purified 70S ribosomes were then resuspended in poly(U) buffer A (20 mM K-HEPES, pH 7.5, 500 mM NH_4_Cl, 10 mM Mg(OAc)_2_) and loaded on a 10 ml poly(U) Sepharose 4B column. The flow through fraction, containing the S1-free 70S ribosomes, was collected, concentrated and dialyzed into dissociation buffer (20 mM K-HEPES, pH 7.5, 200 mM NH_4_Cl, 1 mM Mg(OAc)_2_). The sample was loaded on 15–30% sucrose gradient that was centrifuged in SW28 rotor at 58,000 g for 18 hours to separate 30S and 50S subunits. After the run, the gradient was fractionated, 50S and 30S peak fractions were collected, concentrated, dialyzed against reassociation buffer (20 mM K-HEPES, pH 7.5, 120 mM KOAc, 10 mM NH_4_Cl, 20 mM Mg(OAc)_2_) and were mixed in 1:1 ratio of A_260_ units to have an excess of 30S subunits. The sample was incubated at 40°C for 1 hour followed by 10 minutes on ice and was layered on 15–30% sucrose gradient that was centrifuged in SW28 rotor at 58,000 g for 18 hours to separate excess 30S subunits from reassociated 70S ribosomes. After the run, the gradient was fractionated and 70S peak fractions were collected and concentrated to 20-35 mg/ml. The purified 70S ribosomes were dialyzed into EM buffer (20 mM K-HEPES, pH 7.5, 120 mM KOAc, 10 mM NH_4_Cl, 20 mM Mg(OAc)_2_, 10 µM ZnCl_2_), flash frozen, and stored as small aliquots at −80°C.

### E. coli small ribosomal subunit protein bS1

*E. coli* bS1 containing N-terminal (His)_10__TwinStrep-tag was overexpressed from pAX1_(His)_10_-TwinStrep-HRV3C-rpsA in the *E. coli* LACR II strain. For expression, 6 L of LB culture (50 µg/ml Kanamycin) was induced at an OD_600_ of 0.6-0.8 with 1 mM IPTG for 3 hours at 37°C. Cells were harvested by centrifugation, resuspended in 3 volumes of lysis buffer (20 mM Tris-HCl, pH 7.5, 150 mM NH_4_Cl, 500 mM KCl, 5% glycerol, 0.1 mM PMSF, 1 mM benzamidine, 2 mM β-mercaptoethanol, EDTA-free protease inhibitor cocktail (Sigma-Aldrich cOmplete, 1 tablet/50ml)) and lysed using sonication. The lysate was cleared using a Type 45 Ti rotor (Beckman) at 40,000 g for 30 minutes. After increasing the NH_4_Cl concentration of the supernatant to 1 M to dissociate bS1 from 70S ribosomes the sample was centrifuged in a Type 70 Ti rotor (Beckman) at 60,000 g for 2 hours. The supernatant (containing bS1) was loaded on 5 ml Ni-HiTrap HP column (GE Healthcare) equilibrated with IMAC buffer A (20 mM Tris-HCl, pH 7.5, 1 M NH_4_Cl, 500 mM KCl, 5% glycerol, 0.1 mM PMSF, 1 mM benzamidine, 2 mM β-mercaptoethanol) and after extensive washing with 2% followed by 5% IMAC buffer B (same as IMAC buffer A but 250 mM imidazole), the protein was eluted with 100% IMAC buffer B. Peak fractions containing bS1 were directly loaded on a 5 ml StrepTrap HP column (GE Healthcare) equilibrated with Strep binding buffer (20 mM Tris-HCl, pH 7.5, 40 mM NH_4_Cl, 150 mM KCl, 5% glycerol, 0.1 mM PMSF, 1 mM benzamidine, 2 mM β-mercaptoethanol) and the protein was eluted with Strep elution buffer (20 mM Tris-HCl, pH 7.5, 40 mM NH_4_Cl, 5% glycerol, 0.1 mM PMSF, 1 mM benzamidine, 2 mM β-mercaptoethanol, 5 mM D-Desthiobiotin). The peak fractions were directly loaded on 5 ml HiTrap Q HP column (GE Healthcare) equilibrated with Q buffer A (20 mM Tris-HCl, pH 7.5, 40 mM NH_4_Cl, 5% glycerol, 0.1 mM PMSF, 1 mM benzamidine, 2 mM β-mercaptoethanol) and eluted using a linear gradient of 0-100% Q buffer B (20 mM Tris-HCl, pH 7.5, 40 mM NH_4_Cl, 1 M KCl, 5% glycerol, 0.1 mM PMSF, 1 mM benzamidine, 2 mM β-mercaptoethanol) over 20 column volumes. The sample was dialyzed overnight in the presence of His-tagged HRV3C (PreScission) protease (1 mg HRV3C per 8 mg of protein) into dialysis buffer (20 mM Tris-HCl, pH 7.5, 1 M NH_4_Cl, 500 mM KCl, 5% glycerol, 2 mM β-mercaptoethanol). Uncleaved protein, the cleaved (His)_10_-TwinStrep-tag and HRV3C were selectively removed using the IMAC column; since cleaved bS1 weakly binds to the IMAC column it was eluted with 12% IMAC buffer B. The peak was concentrated and dialyzed into assembly buffer (5 mM HEPES, pH 7.5, 100 mM KOAc, 10 mM Mg(OAc)_2_, 0.5 mM TCEP). The final protein was concentrated to ∼50 mg/ml and aliquots were flash frozen and stored at −20°C.

### E. coli Phenylalanyl-tRNA synthetase (PheRS)

*E. coli* PheRS with an N-terminally (His)_10_-tagged α-subunit was overexpressed from pAX0_(His)_10_-HRV3C-pheS_pheT in the *E. coli* LACR II strain. For expression, 6 L of LB culture (50 µg/ml Kanamycin) were induced at an OD_600_ of 0.6-0.8 with 0.5 mM IPTG for 3 hours at 37°C. Cells were harvested by centrifugation, resuspended in 5 volumes of lysis buffer (50 mM HEPES, pH 7.5, 1 M NH_4_Cl, 10 mM MgCl_2_, 0.1 mM PMSF, 1 mM benzamidine, 2 mM β-mercaptoethanol, DNase I (0.5 mg/250 g cell), EDTA-free protease inhibitor cocktail (Sigma-Aldrich cOmplete, 1 tablet/50 ml)) and lysed using sonication. The lysate was cleared using a Type 45 Ti rotor (Beckman) at 40,000 g for 1 hour. The supernatant was loaded on a 5 ml Ni-HiTrap HP column (GE Healthcare) equilibrated with IMAC buffer A (50 mM HEPES, pH 7.5, 1 M NH_4_Cl, 10 mM MgCl_2_, 10 mM imidazole, 0.1 mM PMSF, 1 mM benzamidine, 2 mM β-mercaptoethanol) and after extensive washing with Buffer A followed by 5% IMAC buffer B (same as IMAC buffer A but 400 mM imidazole) the protein was eluted using a linear gradient of 5-100% IMAC buffer B. Peak fractions were pooled, and dialyzed overnight in the presence of His-tagged HRV3C (PreScission) protease (1 mg HRV3C per 8 mg of protein) into dialysis buffer (50 mM HEPES, pH 7.5, 1 M NH_4_Cl, 10 mM MgCl_2_, 2 mM β-mercaptoethanol). Uncleaved protein, the cleaved (His)_10_-tag and HRV3C were selectively removed using the IMAC column and collecting the flow-through containing cleaved PheRS. The sample was dialyzed into Q binding buffer (10 mM HEPES, pH 7.5, 50 mM NaCl, 1 mM DTT, 0.1 mM PMSF, 1 mM benzamidine) until the conductivity was ≤ 6mS/cm. PheRS was loaded on two 5 ml HiTrap Q columns (GE Healthcare) equilibrated by Q binding buffer and eluted using a linear gradient into Q binding buffer containing 1 M NaCl over 10 column volumes. The peak was concentrated and dialyzed into storage buffer (50 mM HEPES, pH 7.5, 100 mM NaCl, 2 mM DTT). The final protein was concentrated to ∼50 mg/ml and aliquots were flash frozen and stored at −20°C.

### tRNA purification and aminoacylation

The tRNAs were expressed, purified and aminoacylated as was previously described (*31, 32*).

*E. coli* HMS174 cells overexpressing tRNA^fMet^ or tRNA^Phe^ were grown in LB (100 µg/ml Ampicillin) for 24 hours at 37°C. Cells were harvested by centrifugation and resuspended in 10 ml lysis buffer per liter of culture (10 mM Tris-HCl, pH 7.5, 10 mM Mg(OAc)_2_). An equal volume of phenol pH 4.3 was added to the sample and vortexed twice for 30 seconds. The aqueous phase was separated from the organic phase by centrifugation at 27,000 g, 20°C for 30 minutes and was ethanol precipitated by addition of 3 volumes of ethanol. After 1 hour incubation at −20°C the sample was centrifuged at 8,600 g, 4°C for 30 minutes. To separate high molecular weight nucleic acids, the pellet was resuspended in 50 ml 1 M NaCl by vortexing and rolling at room temperature and was cleared by centrifugation at 8,600 g, 4°C for 5 minutes. The supernatant was precipitated by addition of 3 volumes of ethanol and was kept overnight at −20°C. Following centrifugation at 8,600 g, 4°C for 20 minutes, the pellet was resuspended in 25 ml 1.5 M Tris-HCl pH 8.8 and was incubated in a water bath at 37°C for 2 hours in order to deacylate tRNAs. The total tRNA was ethanol precipitated by addition of 3 volumes of ethanol.

*E. coli* tRNA^fMet^ purification: The total tRNA pellet was resuspended in Q-sepharose A buffer (20 mM Tris-HCl, pH 7.5, 8 mM MgCl_2_, 200 mM NaCl, 0.1 mM EDTA). The sample was loaded on a 5 ml HiTrap Q FF column (GE Healthcare) and was eluted using a linear gradient 0-60% into Q-sepharose B buffer (20 mM Tris-HCl, pH 7.5, 8 mM MgCl_2_, 1 M NaCl, 0.1 mM EDTA) over 20 column volumes. Peak fractions were pooled and dialyzed into tRNA storage buffer (10 mM NH_4_OAc, pH 5.0, 50 mM KCl). The tRNA was concentrated to ∼400 µM and aliquots were flash frozen and stored at −80°C.

*E. coli* tRNA^Phe^ and Phe-tRNA^Phe^ purification: The ethanol precipitated total tRNA pellet was resuspended in Phe-sepharose A buffer (20 mM NaOAc, pH 5.3, 10 mM MgCl_2_, 1.5 M (NH_4_)_2_SO_4_) and was loaded on a 50 ml Phenyl Sepharose column (GE Healthcare). After one column volume wash step the tRNAs were eluted with a linear gradient of Phe-sepharose B buffer (20 mM NaOAc, pH 5.3, 10 mM MgCl_2_) 0-60% for 2.3 column volumes, followed by 0.5 column volumes at 60% and 2 column volumes at 100%. Peak fractions with conductivity between 145-110 mS/cm were pooled, the (NH_4_)_2_SO_4_ concentration was adjusted to ≥ 1.7 M, and the sample was loaded on 54 ml TSKgel® Phenyl-5PW column (Tosoh Bioscience) equilibrated wit 5PW buffer A (10 mM NH_4_OAc, pH 6.3, 1.7 M (NH_4_)_2_SO_4_). tRNAs were eluted using a linear gradient of 10-35% 5PW buffer B (10 mM NH_4_OAc, pH 6.3) for 4 column volumes. Peak fractions with conductivity between 176-181 mS/cm were pooled and dialyzed into aminoacylation reaction buffer. After aminoacylation (see next section), Phe-tRNA^Phe^ was purified on Phenyl-5PW column the same way as tRNA^Phe^. It elutes as a single peak at lower conductivity than the deacylated tRNA. Peak fractions were pooled and dialyzed into tRNA storage buffer. The tRNA was concentrated to ∼350 µM and aliquots were flash frozen and stored at −80°C.

tRNA^Phe^ aminoacylation: 20 µM tRNA^Phe^, 200 µM phenylalanine, 4 mM ATP, 0.2 µM PheRS, and 2 U/ml pyrophosphatase (Sigma-Aldrich) were mixed in reaction buffer (20 mM Tris-HCl, pH 7.5, 7 mM MgCl_2_, 150 mM KCl) and incubated for 30 minutes at 37°C. The sample was precipitated by addition of three volumes of ethanol. The pellet was resuspended in 5PW A buffer and loaded on a Phenyl-5PW column (see previous section).

### Oligonucleotide scaffold preparation

DNA (Sigma-Aldrich) and RNA (Dharmacon) oligonucleotides were chemically synthesized and purified by the manufacturer. RNA was deprotected following the protocols provided by the manufacturer. Both DNA and RNA were dissolved in RNase free water and aliquots were stored at −80°C.

For nucleic acid scaffold assembly, template DNA (tDNA) and mRNA were mixed in a 1:1 molar ratio in reconstitution buffer (10 mM HEPES, pH 7.0, 40 mM KOAc, 5 mM Mg(OAc)_2_) and annealed by heating to 95°C followed by stepwise cooling to 10°C in a PCR machine; non-template DNA (ntDNA) was added during complex formation.

### Binding assay

The expressome complex was assembled by mixing *E. coli* 70S ribosomes (2 µM final concentration), nucleic acid scaffold (tDNA, mRNA) and *E. coli* bS1 in low (20 mM HEPES, pH 7.8, 50 mM NaCl, 25 mM MgCl_2_, 20 μM ZnCl_2_, 0.5 mM TCEP) or high salt buffer (20 mM K-HEPES, pH 7.5, 120 mM KOAc, 10 mM NH_4_Cl, 20 mM Mg(OAc)_2_, 10 µM ZnCl_2_, 0.5 mM TCEP) and incubated for 15 min at 37°C. *E. coli* tRNA^fMet^, *E. coli* Phe-tRNA^Phe^ and *E. coli* RNAP were added followed by incubation for 5 min at 37°C after the addition of each component. Finally ntDNA was added and the sample was incubated for 15 min at 37°C. The molar ratios of the components were the following: 70S:S1:nucleic acid scaffold: tRNA^fMet^:Phe-tRNA^Phe^:RNAP=1:1.5:4.5:2:2:4. 30 µl reaction mixtures were layered on top of 15-30% sucrose gradient and were centrifuged at 43,500 g for 16 hours at 4°C, in an SW60 rotor (Beckman). Sucrose solutions for gradient preparation were prepared either in low or high salt buffer. The gradient was fractionated from top to bottom in 150 µl fractions. Peak fractions (OD_260_) were combined and concentrated. Samples were separated on Nu-PAGE™ 4-12% Bis-Tris gel (Invitrogen) and stained with Coomassie Blue G-250.

### Sample preparation for cryo-EM analysis

Expressome complexes RNA-34 and RNA-38 with or without NusG saturation were purified by sucrose gradient centrifugation: The expressome complex was assembled by mixing *E. coli* 70S ribosomes (2 µM final concentration), nucleic acid scaffold (tDNA, mRNA) and *E. coli* bS1 in assembly buffer (5 mM HEPES, pH 7.5, 100 mM KOAc, 10 mM Mg(OAc)_2_, 0.5 mM TCEP) and incubated for 15 min at 37°C. *E. coli* tRNA^fMet^, *E. coli* Phe-tRNA^Phe^, *E. coli* RNAP and ntDNA were added followed by incubation for 5 min at 37°C after the addition of each component. *E. coli* NusG was added and the sample was incubated for 30 min at 37°C. The molar ratios of the components were the following: 70S:bS1:nucleic acid scaffold: tRNA^fMet^:Phe-tRNA^Phe^:RNAP:NusG=1:4:4:2:2:10:12.5. Bis(sulfosuccinimidyl)suberate (BS3) was added at 5 mM final concentration for cross-linking and the sample was incubated on ice for one hour. Excess proteins, tRNAs and nucleic acid scaffold were removed by layering 100 µl of sample on top of four 15-30% sucrose gradients and centrifugation at 43,500 g for 18 hours at 4°C, in an SW60 rotor (Beckman). The sucrose solution was prepared with EM buffer containing 10 mM NH_4_Cl that stopped the cross-linking reaction. The gradient was fractionated from bottom to top in 150 µl fractions. Peak fractions (OD_260_) were combined, concentrated, and the sample was dialyzed overnight against EM buffer containing 0.5 mM TCEP. The final concentration, measured in absorbance units at 260nm, was between 0.14-0.24 µM (OD_260_ 6-10) before grid freezing. Optionally, additional NusG was added at 25 µM final concentration to saturate NusG occupancy.

### Cryo-EM grid preparation and data collection

Quantifoil R2/2 300 mesh holey carbon copper grids (Quantifoil Micro Tools, Großlöbichau, Germany) were glow-discharged (ELMO Glow Discharge System) for 30 s at 2.5 mA prior to the application of 3.5 µl sample and plunge-frozen in liquid ethane using a Vitrobot Mark IV (FEI) with 95% chamber humidity at 10°C. The grids were imaged using a 300 keV Titan KRIOS (FEI) with a K2 Summit direct electron detector (Gatan) at pixel size of 1.052 Å/px (RNA-38 data) or 1.075 Å/px (RNA-34 data). Movies with 40-41 frames were collected with a total electron dose of 42-51 e^−^/Å^2^ at a rate of 6.4-7.3 e^−^/Å^2^/sec in counting mode with defocus values in the range −0.7 to −3.5 µm (Table S1).

### Cryo-EM data processing

Image frames were aligned and averaged with MotionCor2 (*33*), and contrast transfer function (CTF) parameters were calculated with Gctf (*34*). All subsequent steps were performed in RELION-3 (*35*). Automated particle picking was perfomed using templates generated from the two-dimensional class averages of 2000 manually selected particles. Particles not containing ribosomes were discarded following reference-free two-dimensional classification. Maps for three-dimensional references were obtained using the initial model tool in RELION-3, and an initial round of three-dimensional classification was performed to further remove particles that were poorly aligned. The remaining number of particle images were 546512, 387633, 272809, 157304 for the samples RNA-38, RNA-38+NusG, RNA-34 and RNA-34+NusG respectively.

For the uncoupled expressome data (RNA-38), separation of expressome particles from ribosomes through conventional 3D classification was ineffective, likely due to the heterogeneity in RNAP position. As an alternative, particles were first extracted with re-centering on the partially-occupied RNAP density. A map was produced from these particles using a mask around the full expressome. The signal corresponding to the ribosome was subtracted on a per-particle basis. Without the stronger signal from the ribosome, particles could be separated into those that contained RNAP and those that did not by two-dimensional classification with a mask of 160 Å. The option ‘ignore CTFs until first peak’ was found to significantly improve the outcome of this process. Signal subtraction was reverted once the expressome particle subset was obtained. The success of this approach in removing particles without RNAP was evident in the substantially improved signal for RNAP (Fig S1E). The final number of expressome particle images was 32195. Relative orientation analysis and further particle selection was performed to isolate clusters of preferred molecular states (see below).

For the coupled expressome data (RNA-38+NusG), 3D classification without alignment was performed, a mask that included the partially-occupied RNAP density and neighboring ribosome surface was applied and a resolution limit of 20 Å (Fig S4A). RNAP-containing particles were selected and 3D classification was repeated, leading to identification of 15327 particles with well-resolved features for both ribosome and RNAP. Multibody refinement was performed with masks around the ribosome and RNAP (*23*). Masks were created by manually erasing density from the consensus map and low-pass filtering to 30 Å. Exclusion of the RNAP β′-ZF domain, which inserts into the ribosome, from the masked area was necessary for accurate alignment.

For the collided expressome data (RNA-34+NusG), the map of the full expressome was obtained from 5360 particles selected following masked 3D classification without alignment (Fig S7B). High-resolution maps of the ribosome and RNAP were obtained from 45774 particles selected by ribosome signal-subtracted, as described for the uncoupled expressome. For the collided expressome lacking NusG, 18552 particles were selected by ribosome signal-subtraction.

Ribosome maps were further improved by focused refinement with masks around either the 50S, 30S-head or 30S-body (Fig S2A). While the nominal resolution was modestly improved (∼0.1 Å), surface regions of initially lower resolution were improved significantly. Composite maps were generated with the Phenix routine combine-focused-maps (*36*).

### Model building

We constructed initial models of the different complexes by combining the X-ray structure of the empty *E. coli* 70S ribosome (PDB ID 4YBB) (*37*), and the EM structure of an *E. coli* RNAP elongation complex (PDB ID 6ALH) (*38*). tRNA models were derived from crystal structures (tRNA^fMet^: PDB ID 2FMT, and Phe-tRNA^Phe^: PDB ID 3L0U) (*39, 40*). The individual models were placed into the EM maps using UCSF Chimera (*41*) followed by rigid body refinement in Phenix (*36*). High resolution X-ray and EM structures of the *Thermus thermophilus* and *E. coli* ribosome were used to guide model building (*42-44*). The mRNA was built *de novo*, and all models were adjusted to fit the EM maps in Coot (*45*). Two *E. coli* ribosome regions are modeled substantially differently between previous structures: the C-terminus of bS21, and the N-terminus of uS19. We found that our map clearly favored a model similar to only one reported structure (PDB ID 5MDV) (*46*) (Fig S2D). This was followed by iterative rounds of real-space refinement using secondary structure restraints and geometry optimization in Phenix (*36*), manual inspection, and model adjustments in Coot.

The accession numbers for the four cryo-EM reconstructions (uncoupled expressome without saturating NusG and with RNA-38 mRNA, coupled expressome with saturating NusG and with RNA-38 mRNA, collided expressome without saturating NusG and with RNA-34 mRNA, collided expressome with saturating NusG and with RNA-34 mRNA) reported in this paper are … Fitted models were deposited in the PDB under accession numbers WWWW (uncoupled expressome without saturating NusG and with RNA-38 mRNA), XXXX (coupled expressome with saturating NusG and with RNA-38 mRNA), YYYY (collided expressome without saturating NusG and with RNA-34 mRNA), ZZZZ (collided expressome with saturating NusG and with RNA-34 mRNA).

### Quantification of expressome relative orientations and particle subset selection

Reconstructed maps of the ribosome and RNAP were first re-oriented so that the Z-axis was aligned and centered on the mRNA entrance-channel and exit-channel respectively. Following low-pass filtering to 30 Å, these maps were used as initial models for 3D refinement. This yielded data for the Euler angle assignment of particle images to maps in which the first angle (rot) represents rotation about the mRNA axis, and the following angles (tilt, psi) represent orthogonal rotations. During reconstructing RNAP and ribosome maps, particle images were re-extracted from micrographs to permit re-centering. Information for which RNAP and ribosome image pairs correspond to a shared expressome complex were maintained in the RELION data file within the entry ‘ImageOriginalName’.

For each expressome complex, six Euler angles were thereby obtained: ribosome-rot, ribosome-tilt, ribosome-psi, RNAP-rot, RNAP-tilt and RNAP-psi. Rotation matrices were derived for ribosome and RNAP using each set of Euler angles (*47*). The rotational position of RNAP relative to the ribosome was calculated as the product of the rotation matrix for RNAP and the inverse rotation matrix for the ribosome. The resulting matrix was converted into Euler angles in the sequence Z_1_Y_2_X_3_ to allow visual representation and interpretation. The first angle (Z) describes rotation of RNAP about the mRNA axis. Plots of per-particle relative orientation were generated with gnuplot.

For the uncoupled expressome data, particles within clusters of shared relative orientation were selected on the basis of thresholds for all the three Euler angles. The values for thresholds were selected to obtain 1300-1500 particles for each cluster: this was empirically determined to balance the improved particle homogeneity permitted by tighter thresholds with the signal-to-noise limitations resulting from lower particle number. Of the seven defined clusters, only three (1, 2 and 6) contained particles that were sufficiently homogeneous to produce a map in which RNAP coordinates could be docked automatically and unambiguously with UCSF Chimera (*41*). Angles measured for the RNAP orientation assigned were in close agreement with that predicted as the center of the cluster (Fig S3A), validating the approach. For the remaining clusters (3, 4, 5 and off-axis), values at the approximate center of the cluster were applied to rotate RNAP coordinates, while translation was determined by fitting into maps without rotation. Improved RNAP maps were obtained for clusters 1, 2, 6 by multibody refinement (*23*), followed by extraction of ribosome-subtracted particle images, and 3D refinement (Fig S3D). Structural overlay with RNAP-NusG complex coordinates (PDB ID 6C6U) (*13*) confirmed the presence of NusG in the map for cluster 6.

### Negative stain grid preparation and data collection

Samples were prepared with purified *E. coli* components and all contained 70S ribosomes (200 nM), Phe-tRNA^Phe^ (400 nM), tRNA^fMet^ (400 nM) and RNAP (2 µM) in EM buffer. Samples saturated with bS1 were prepared by addition of purified bS1 to a final concentration of 1 µM before addition of nucleic acid scaffold or tRNAs. For samples with nucleic acid scaffold, annealed tDNA-RNA was added to final concentration of 4 µM before addition of tRNAs, and ntDNA was added to the same concentration after addition of RNAP. Samples were diluted 40-fold in EM buffer to final concentrations of 5 nM ribosome and 50 nM RNAP and applied to grids without further purification.

Grids coated with thin carbon (CF300-CU-50, purchased from Electron Microscopy Sciences) were glow-discharged for 30 s before deposition of sample. Following blotting of excess solution, grids were stained with filtered uranyl acetate solution (1.5% w/v) for 30 s, before blotting again. Images were collected on a Tecnai F20 transmission electron microscope at 200 keV with a Gatan CCD detector with settings of 3.4 Å/px pixel size, −0.7 µm defocus and 25 electrons/Å2 dose. For each dataset, approximately 25 000 ribosome-containing particles were selected from approximately 1000 micrographs. Images were extracted (box size 425 Å), and masks of 300-400 Å were applied during two-dimensional classification with RELION-3 software (*35*). Classes representing RNAP:ribosome complexes were identified by comparison to corresponding views of particles containing ribosome only.

### Measurement of clashes in theoretical expressome models

Atomic coordinates for the ribosome and RNAP were derived from the <u>RNA-</u>34+NusG dataset. To determine whether further rotation of RNAP is prohibited by steric clash with the ribosome, a series of RNAP coordinates were generated by rotation about an axis orthogonal to the mRNA axis (axis Y or X) in increments of 2°. From each of these models, a series of coordinates were generated by rotation about the mRNA axis (axis Z) in increments of 2°. Rotation was performed about the mRNA emerging from RNAP, so that all models have the same mRNA pathlength to the ribosome mRNA entrance channel. Clashes between RNAP and the ribosome were measured with PyMOL and defined as separation of less than 2.5 Å between backbone atoms. Due to their flexibility, omega residues 77-91 and NusG residues 46-62 were excluded from the analysis. Angles were normalized to the coordinate system used for relative orientation analysis to allow comparison.

To relate RNAP position to the shortest sterically-allowed mRNA pathlength, a series of RNAP coordinates were first generated by translation of RNAP along the mRNA axis in increments of 0.5 Å with a range of −20 to +29 Å, where the positive direction represents displacement away from the ribosome. mRNA path lengths were measured between phosphate of C27 on the ribosome to A40 on RNAP. From each of these 99 models, 180 coordinates were generated by rotation of RNAP about the mRNA axis in increments of 2°, yielding a total of 17,820 coordinates. Clashes between RNAP and the ribosome were measured as described above. Models were defined as disallowed if greater than 5 clashes were detected. The calculation was repeated with models that were translated orthogonal to the mRNA axis (4 Å) or tilted (5°) to verify that conclusions were insensitive to these changes. The number of particles in each rotational increment of RNAP about the mRNA axis was calculated from the relative orientation analysis.

## Acknowledgments

We thank Julio Ortiz, Corinne Crucifix, Xieyang Guo and Tat Cheung Cheng for help with data collection at the IGBMC. We thank Wim Hagen and Felix Weis for help with data collection at the EMBL in Heidelberg, Germany. We thank Venki Ramakrishnan and Ann C. Kelley for their valuable help in ribosome and tRNA purification. This work has been supported by iNEXT PID 6979, funded by the Horizon 2020 program of the European Union. We acknowledge the European Synchrotron Radiation Facility for the provision of microscope time on CM01, and we thank Gregory Effantin and Michael Hons for their assistance. We thank members of the Weixlbaumer lab for critical reading of the manuscript.

## Author contributions

M.W.W. and M.T. performed experiments, binding assays, electron microscopy, and data analysis. C.Z. participated in data analysis and purification of NusG. M.W.W., M.T., V.V., A.D.E., M.A. A.W. built and refined atomic models. A.W. designed and supervised research. M.W.W., M.T., and A.W. prepared the manuscript with input from all authors. Authors declare no competing interests.

## Data and materials availability

Electron density maps for uncoupled, coupled, and collided expressomes (with and without NusG) were deposited in the EM database (EMD-WWWW, EMD-XXXX, EMD-YYYY, EMD-ZZZZ). Refined coordinates were deposited in the PDB database under accession codes WWWW (uncoupled expressome), XXXX (coupled expressome), YYYY (collided expressome with NusG), and ZZZZ (collided expressome without NusG).

## References and Notes

1. R. Byrne, J. G. Levin, H. A. Bladen, M. W. Nirenberg, The *in vitro* formation of a DNA-ribosome complex. Proc Natl Acad Sci USA. 52, 140–148 (1964).

2. O. L. Miller, B. A. Hamkalo, C. A. Thomas, Visualization of bacterial genes in action. Science. 169, 392–395 (1970).

3. C. Yanofsky, Attenuation in the control of expression of bacterial operons. Nature. 289, 751–758 (1981).

4. J. P. Richardson, Preventing the synthesis of unused transcripts by Rho factor. Cell. 64, 1047–1049 (1991).

5. S. Proshkin, A. R. Rahmouni, A. Mironov, E. Nudler, Cooperation between translating ribosomes and RNA polymerase in transcription elongation. Science. 328, 504–508 (2010).

6. M. Zhu, M. Mori, T. Hwa, X. Dai, Disruption of transcription-translation coordination in Escherichia coli leads to premature transcriptional termination. Nat Microbiol. 54, 1–10 (2019).

7. B. M. Burmann et al., A NusE:NusG Complex Links Transcription and Translation. Science. 328, 501–504 (2010).

8. S. Saxena et al., Escherichia coli transcription factor NusG binds to 70S ribosomes. Mol Microbiol. 108, 495–504 (2018).

9. R. Kohler, R. A. Mooney, D. J. Mills, R. Landick, P. Cramer, Architecture of a transcribing-translating expressome. Science. 356, 194–197 (2017).

10. G. Demo et al., Structure of RNA polymerase bound to ribosomal 30S subunit. Elife. 6, 94 (2017).

11. H. Fan et al., Transcription-translation coupling: direct interactions of RNA polymerase with ribosomes and ribosomal subunits. Nucleic Acids Res. 45, 11043–11055 (2017).

12. D. Castro-Roa, N. Zenkin, In vitro experimental system for analysis of transcription-translation coupling. Nucleic Acids Res. 40, e45–e45 (2012).

13. J. Y. Kang et al., Structural Basis for Transcript Elongation Control by NusG Family Universal Regulators. Cell. 173, 1650–1662.e14 (2018).

14. S. L. Sullivan, M. E. Gottesman, Requirement for E. coli NusG protein in factor-dependent transcription termination. Cell. 68, 989–994 (1992).

15. M. R. Lawson et al., Mechanism for the Regulated Control of Bacterial Transcription Termination by a Universal Adaptor Protein. Mol Cell. 71, 911–922.e4 (2018).

16. M. Turtola, G. A. Belogurov, NusG inhibits RNA polymerase backtracking by stabilizing the minimal transcription bubble. Elife. 5 (2016), doi:10.7554/eLife.18096.

17. G. Vauquelin, S. J. Charlton, Exploring avidity: understanding the potential gains in functional affinity and target residence time of bivalent and heterobivalent ligands. Br. J. Pharmacol. 168, 1771–1785 (2013).

18. S. Takyar, R. P. Hickerson, H. F. Noller, mRNA helicase activity of the ribosome. Cell. 120, 49–58 (2005).

19. H. Amiri, H. F. Noller, Structural evidence for product stabilization by the ribosomal mRNA helicase. RNA. 25, 364–375 (2019).

20. X. Qu et al., The ribosome uses two active mechanisms to unwind messenger RNA during translation. Nature. 475, 118–121 (2011).

21. C. L. Chan, R. Landick, The Salmonella typhimurium his operon leader region contains an RNA hairpin-dependent transcription pause site. Mechanistic implications of the effect on pausing of altered RNA hairpins. J Biol Chem. 264, 20796–20804 (1989).

22. I. Gusarov, E. Nudler, The mechanism of intrinsic transcription termination. Mol Cell. 3, 495–504 (1999).

23. T. Nakane, D. Kimanius, E. Lindahl, S. H. Scheres, Characterisation of molecular motions in cryo-EM single-particle data by multi-body refinement in RELION. Elife. 7, 1485 (2018).

24. K. R. Andersen, N. C. Leksa, T. U. Schwartz, Optimized E. coli expression strain LOBSTR eliminates common contaminants from His-tag purification. Proteins. 81, 1857–1861 (2013).

25. K.-A. F. Twist et al., A novel method for the production of in vivo-assembled, recombinant Escherichia coli RNA polymerase lacking the α C-terminal domain. Protein Sci. 20, 986–995 (2011).

26. X. Guo et al., Structural Basis for NusA Stabilized Transcriptional Pausing. Mol Cell. 69, 816–827.e4 (2018).

27. M. N. Vassylyeva et al., Purification, crystallization and initial crystallographic analysis of RNA polymerase holoenzyme from Thermus thermophilus. Acta Crystallogr D Biol Crystallogr. 58, 1497–1500 (2002).

28. D. Moazed, H. F. Noller, Transfer RNA shields specific nucleotides in 16S ribosomal RNA from attack by chemical probes. Cell. 47, 985–994 (1986).

29. G. Blaha et al., Preparation of functional ribosomal complexes and effect of buffer conditions on tRNA positions observed by cryoelectron microscopy. Meth Enzymol. 317, 292–309 (2000).

30. A. R. Subramanian, Structure and functions of ribosomal protein S1. Prog. Nucleic Acid Res. Mol. Biol. 28, 101–142 (1983).

31. R. Jünemann et al., In vivo deuteration of transfer RNAs: overexpression and large-scale purification of deuterated specific tRNAs. Nucleic Acids Res. 24, 907–913 (1996).

32. E. Cayama et al., New chromatographic and biochemical strategies for quick preparative isolation of tRNA. Nucleic Acids Res. 28, E64 (2000).

33. S. Q. Zheng et al., MotionCor2: anisotropic correction of beam-induced motion for improved cryo-electron microscopy. Nat Methods. 14, 331–332 (2017).

34. K. Zhang, Gctf: Real-time CTF determination and correction. J. Struct. Biol. 193, 1–12 (2016).

35. J. Zivanov et al., New tools for automated high-resolution cryo-EM structure determination in RELION-3. Elife. 7 (2018), doi:10.7554/eLife.42166.

36. D. Liebschner et al., Macromolecular structure determination using X-rays, neutrons and electrons: recent developments in Phenix. Acta Crystallogr D Struct Biol. 75, 861–877 (2019).

37. J. Noeske et al., High-resolution structure of the Escherichia coli ribosome. Nat Struct Mol Biol. 22, 336–341 (2015).

38. J. Y. Kang et al., Structural basis of transcription arrest by coliphage HK022 nun in an Escherichia coli RNA polymerase elongation complex. Elife. 6 (2017), doi:10.7554/eLife.25478.

39. E. Schmitt, M. Panvert, S. Blanquet, Y. Mechulam, Crystal structure of methionyl-tRNAfMet transformylase complexed with the initiator formyl-methionyl-tRNAfMet. EMBO J. 17, 6819–6826 (1998).

40. R. T. Byrne, A. L. Konevega, M. V. Rodnina, A. A. Antson, The crystal structure of unmodified tRNAPhe from Escherichia coli. Nucleic Acids Res. 38, 4154–4162 (2010).

41. E. F. Pettersen et al., UCSF Chimera--a visualization system for exploratory research and analysis. J Comput Chem. 25, 1605–1612 (2004).

42. M. Selmer et al., Structure of the 70S ribosome complexed with mRNA and tRNA. Science. 313, 1935–1942 (2006).

43. R. M. Voorhees, A. Weixlbaumer, D. Loakes, A. C. Kelley, V. Ramakrishnan, Insights into substrate stabilization from snapshots of the peptidyl transferase center of the intact 70S ribosome. Nat Struct Mol Biol. 16, 528–533 (2009).

44. N. Fischer et al., Structure of the E. coli ribosome-EF-Tu complex at <3 Å resolution by Cs-corrected cryo-EM. Nature. 520, 567–570 (2015).

45. P. Emsley, K. Cowtan, Coot: model-building tools for molecular graphics. Acta Crystallogr D Biol Crystallogr. 60, 2126–2132 (2004).

46. N. R. James, A. Brown, Y. Gordiyenko, V. Ramakrishnan, Translational termination without a stop codon. Science. 354, 1437–1440 (2016).

47. J. B. Heymann, M. Chagoyen, D. M. Belnap, Common conventions for interchange and archiving of three-dimensional electron microscopy information in structural biology. J. Struct. Biol. 151, 196–207 (2005).

48. Y. Zhang, S. Hong, A. Ruangprasert, G. Skiniotis, C. M. Dunham, Alternative Mode of E-Site tRNA Binding in the Presence of a Downstream mRNA Stem Loop at the Entrance Channel. Structure. 26, 437–445.e3 (2018).

49. A. B. Loveland, A. A. Korostelev, Structural dynamics of protein S1 on the 70S ribosome visualized by ensemble cryo-EM. Methods. 137, 55–66 (2018).

50. M. A. Lauber, J. Rappsilber, J. P. Reilly, Dynamics of ribosomal protein S1 on a bacterial ribosome with cross-linking and mass spectrometry. Mol. Cell Proteomics. 11, 1965–1976 (2012).

