## Supplementary Materials for "Structural basis of transcription-translation coupling and collision in bacteria"

**This PDF file includes:**

Figs. S1 to S11
Tables S1
Captions for Movies S1 to S3

**Other Supplementary Materials for this manuscript include the following:**

Movies S1 to S3

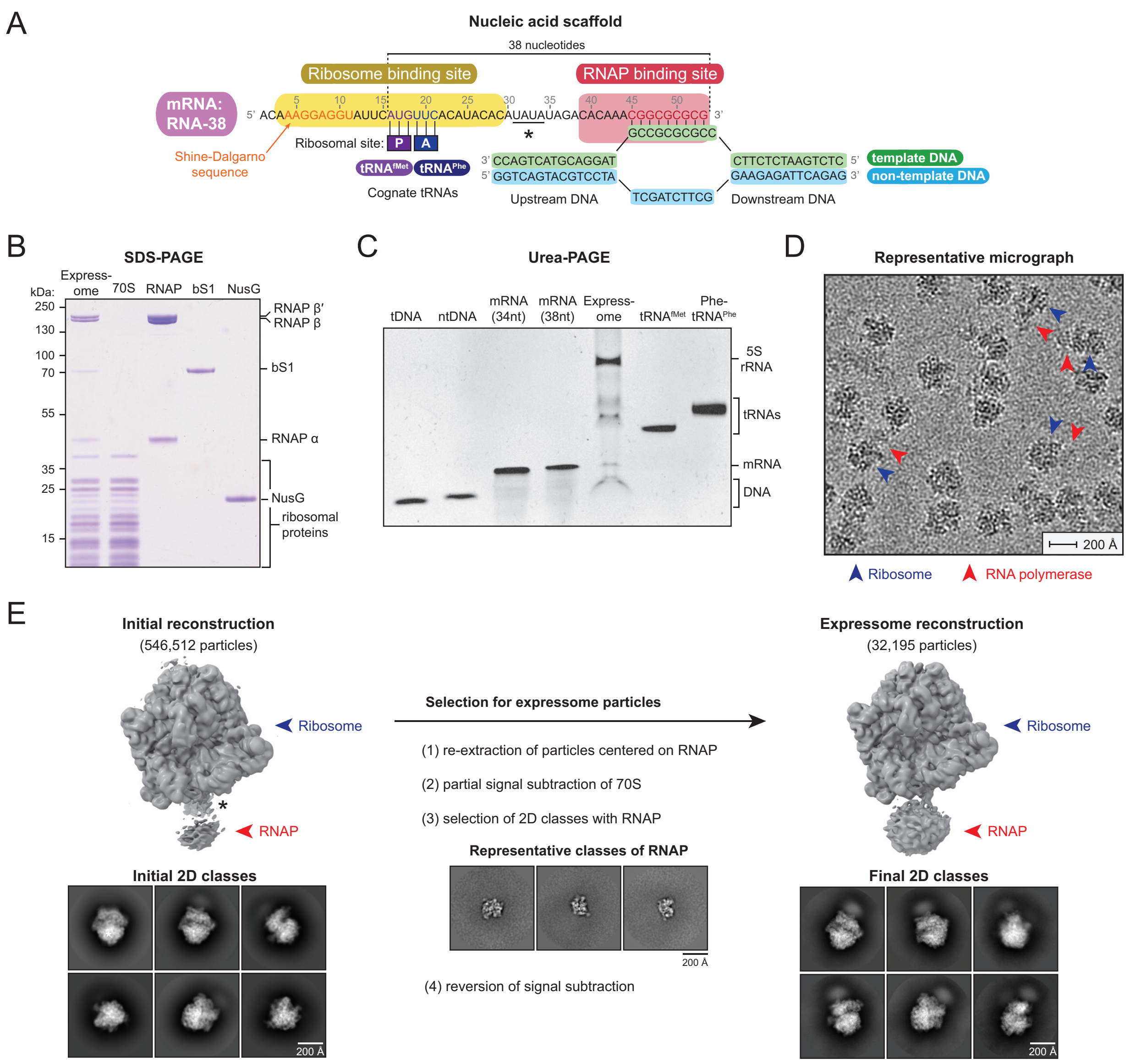


### Figure S1

**Figure S1:** Cryo-EM of the uncoupled expressome. **(A)** Schematic of nucleic acid scaffold used to assemble expressome complexes for cryo-EM. Key sequence features of the mRNA (RNA-38) are colored: Shine-Dalgarno sequence for ribosome binding (orange), P- and A-site codons for tRNA binding (purple and blue), sequence complementary to template DNA (red). Nucleotides removed in the shortened mRNA (RNA-34) are indicated with an asterisk. mRNAs are named according to the number of nucleotides between, and including, the ribosomal P-site and the RNA-DNA hybrid. **(B)** Coomassie-stained SDS-PAGE showing purified *E. coli* protein components used for the assembly of all expressome samples. **(C)** Denaturing urea-PAGE of chemically synthesized (template DNA, tDNA; non-template DNA, ntDNA, mRNA) or *E. coli* nucleic acid components (tRNAs, rRNAs) used for the assembly of all expressome samples (note that presence of sucrose affects migration in the expressome lane). **(D)** Representative cryo electron micrograph of expressomes used for 3D reconstructions. Pairs of red and blue arrows indicate example expressome particles. **(E)** Particle classification scheme based on partial signal subtraction. Initial 3D models and 2D classes showed weak signal corresponding to RNAP, suggestive of partial occupancy (left). Following particle re-centering and subtraction of ribosome signal (see methods), 2D classes with high-resolution RNAP features were selected (center, representative unmasked classes shown). Improved signal for RNAP in both 3D model and 2D classes confirmed selection of expressomes (right). This method also eliminated particles with RNAP bound in an alternative site (marked *). In subsequent samples, ribosomes were saturated with an excess of recombinant ribosomal protein bS1, which eliminated binding of RNAP at this secondary position.

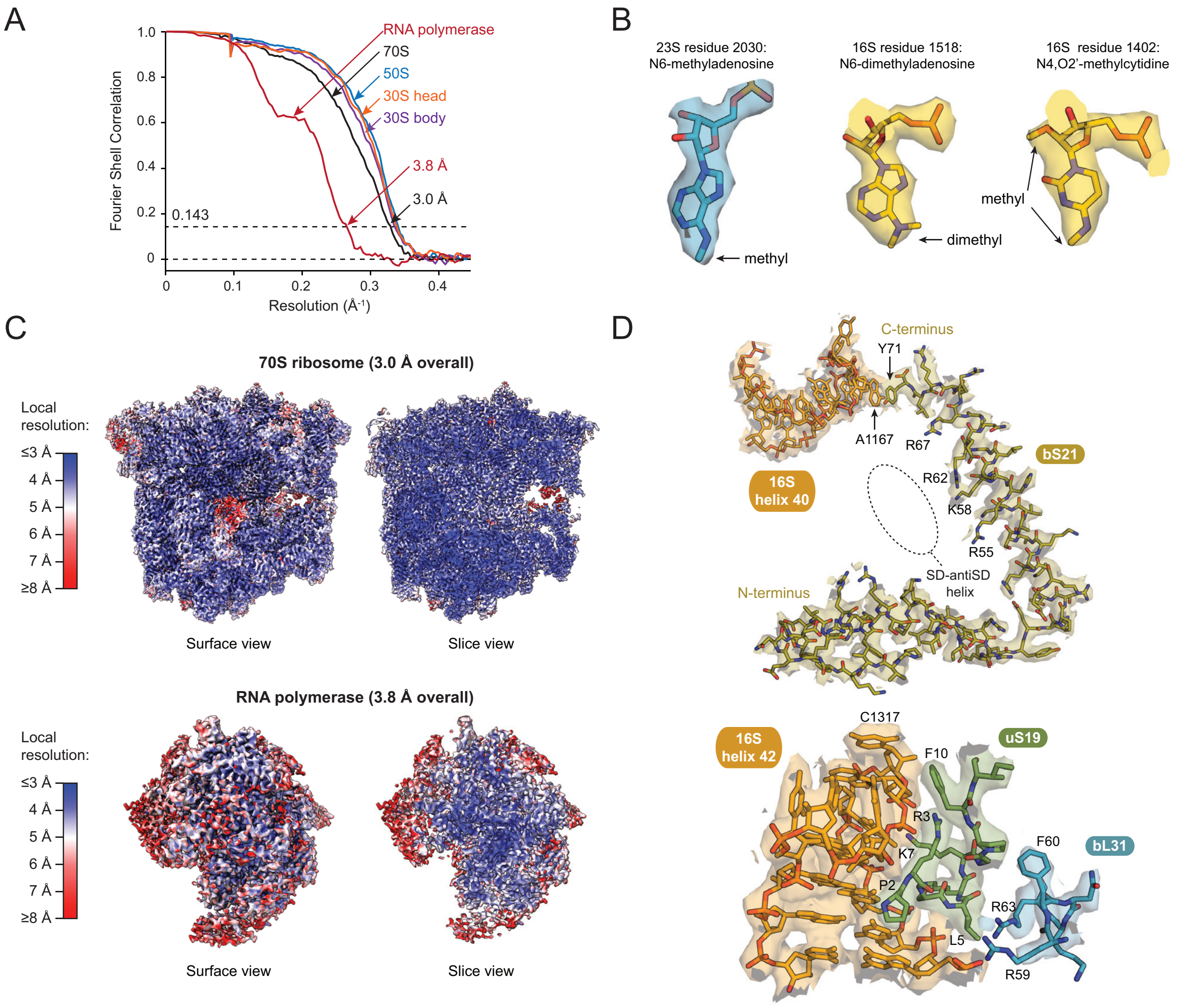


**Figure S2**

**Figure S2:** Resolution and map features of the uncoupled expressome. **(A)** Fourier shell correlation plots for gold standard refinements. Labels indicate the expressome region within the mask used for each refinement. **(B)** Examples of cryo-EM map density for modified nucleic acids indicate map quality. **(C)** Local resolution map of the ribosome and RNAP regions within the uncoupled expressome. The map refined with a mask around the 70S ribosome ranges from ~3 Å in the core to ~4 Å for regions on the surface. Maps produced by focused refinement were more uniform and improved resolution (see methods). The map refined with a mask around RNAP ranges from ~3.8 Å in the core to ~8 Å for domains typically identified as mobile. **(D)** Cryo-EM maps and superimposed models of ribosome regions modeled differently in previous structures (*37*, *44*, *46*). In bS21, the C-terminal residue Y71 stacks against 16S nucleotide

A1167, presenting a series of basic residues (R55, K58, R62, R67) into the solvent-exposed cavity containing the Shine-Dalgarno helix (top panel). The N-terminus of uS19 folds in a cavity of 16S helix 42 and forms an interface for the bL31 C-terminal helix (bottom panel).

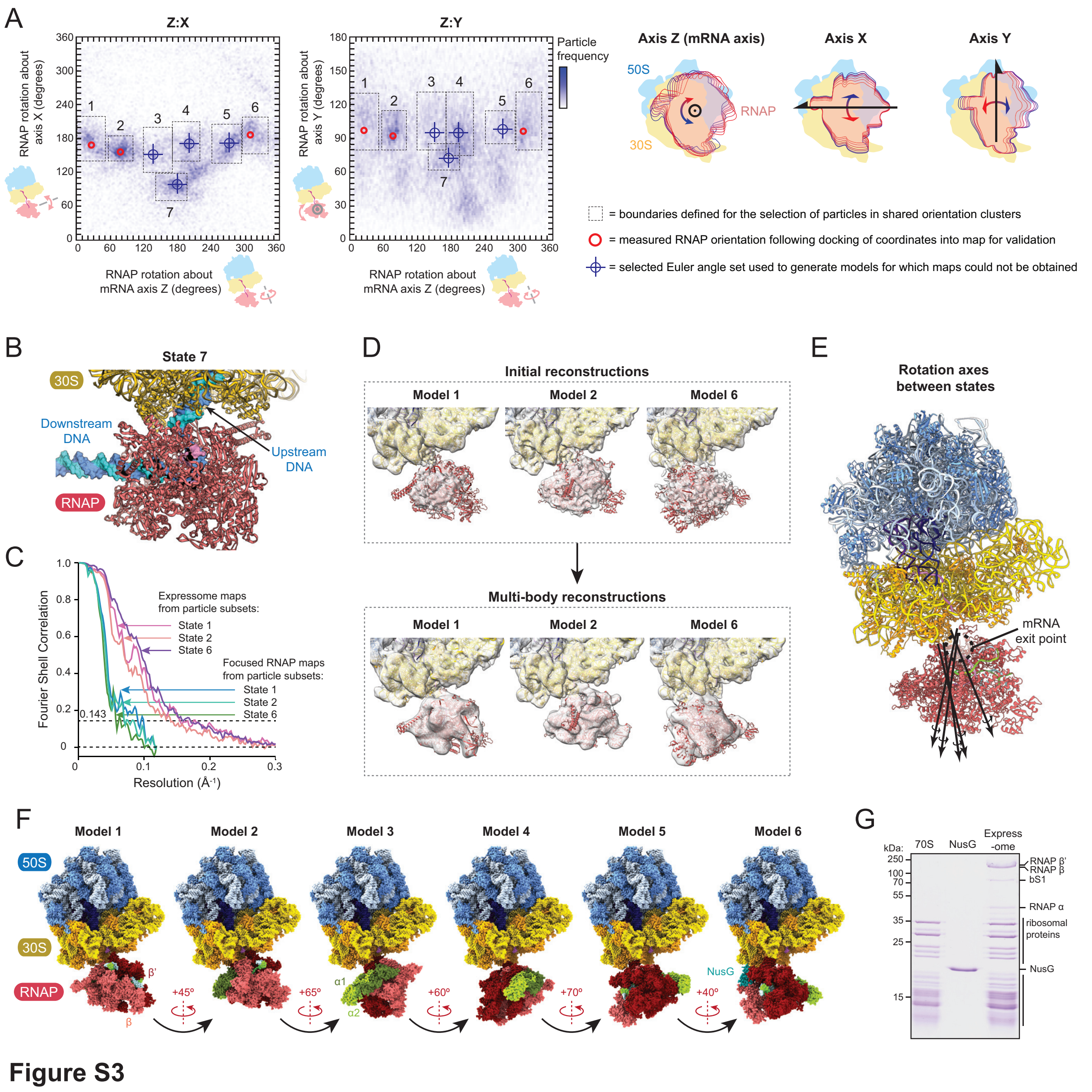


**Figure S3:** Dynamics of the uncoupled expressome. **(A)** Heatmaps of the orientation of RNAP relative to the ribosome in the uncoupled expressome. The spatial definition of the three Euler angles describing relative orientation are shown as cartoons (top). The horizontal axis of both plots indicates rotation of RNAP about an axis connecting the mRNA entrance- and exit-channels in a plane approximately parallel to the surface of the ribosome. The vertical axes of the two plots represent rotation of RNAP about each of the orthogonal axes (X and Y). The plots thereby represent two-dimensional projections of the three-dimensional space describing relative orientation. Shades of blue indicate the frequency of particles in each 2˚ by 2˚ pixel. Dotted lines indicate the boundaries defined for the selection of particles in each of the clusters (marked 1-7). Red circles indicate the measured orientation of RNAP following docking of coordinates into maps produced from the corresponding particle subset. Blue targets indicate the RNAP orientation defined in instances where expressome maps of sufficient resolution to unambiguously dock RNAP could not be obtained. **(B)** Modeled RNAP position of particles in cluster 7 (off-axis) with extended DNA path modeled. The upstream DNA emerging from RNAP would clash with the ribosome making it unlikely to occur in a physiological context. The imaged sample contained 15 base-pairs of upstream DNA. **(C)** Fourier shell correlation plots for expressome maps obtained from particle subsets 1, 2 and 6, and for RNAP maps from subsequent multi-body refinement. Resolutions for expressome maps were between 5.8 Å and 7.2 Å resolution, and RNAP maps were between 10 Å and 15 Å resolution. **(D)** Cryo-EM maps filtered to 8 Å and fitted models for reconstructions obtained from particle subsets in clusters 1, 2 and 6. While limited in resolution, the initial reconstructions (left) displayed sufficient features for automatic docking of ribosome and RNAP. Multi-body refinement with separate masks around the ribosome and RNAP yielded maps with improved features (right) that permitted identification of NusG in model 6 but not model 1 or 2. **(E)** Rotation axes for sequential rotation between states 1 through 6 are approximately perpendicular to the surface of the ribosomes and parallel to the mRNA axis. **(F)** Sphere representation of expressome models for states 1-6 identified by analysis of relative orientation. The approximate angle of RNAP rotation that separates each state is indicated below. The predicted position of NusG in state 6 is indicated. **(G)** NusG co-purifies with expressomes in sub-stoichiometric quantities. Samples saturated with NusG were subsequently prepared by adding excess NusG after gradient purification.

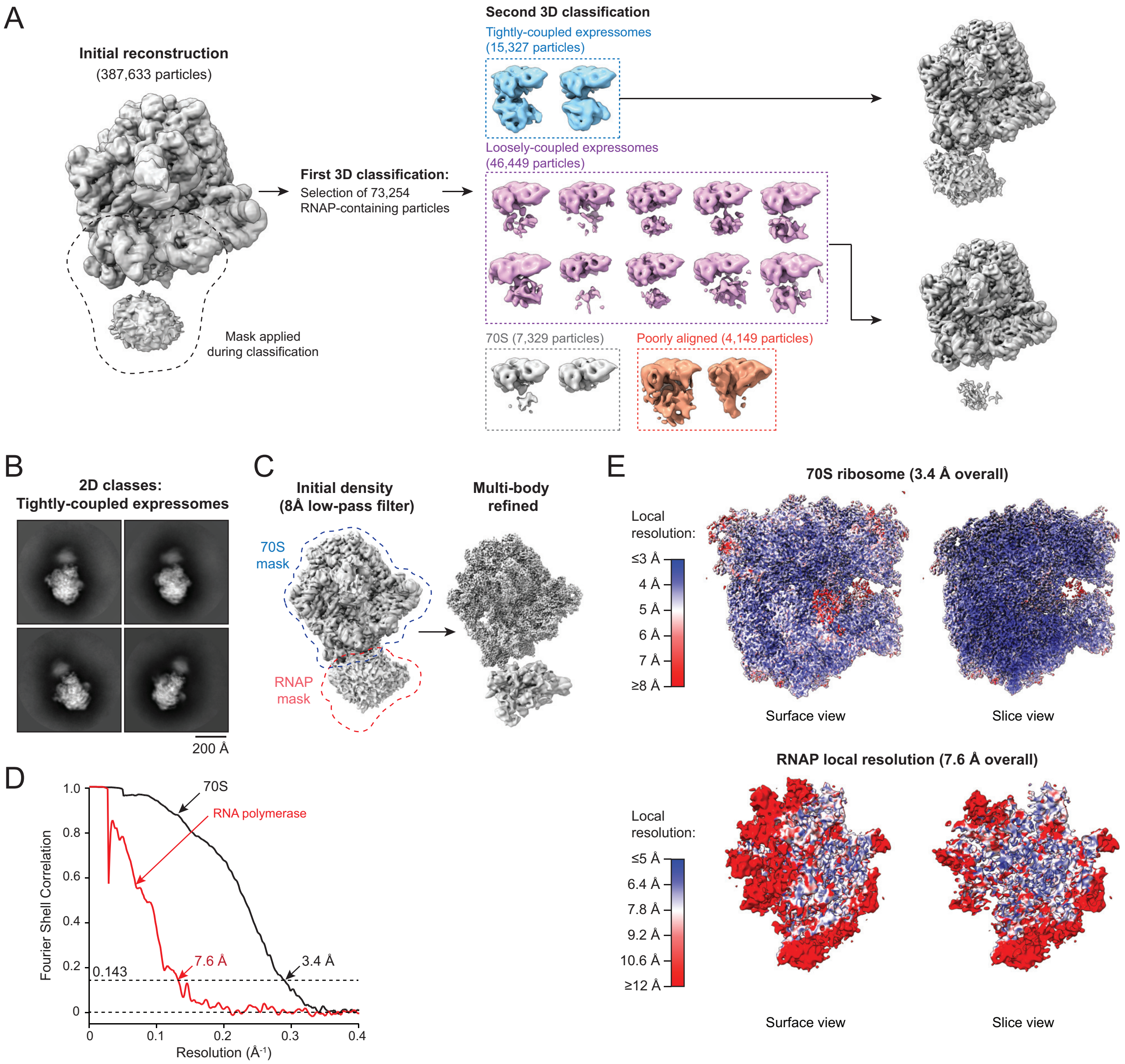


**Figure S4**

**Figure S4:** Cryo-EM of the NusG-coupled expressome. **(A)** 3D classification scheme: Initial 2D and 3D classification produced 387,633 particles which were a mixture of expressomes and ribosomes. To select for expressome particles, two further rounds of 3D classification were performed with a mask around the RNAP region and part of the ribosomal 30S subunit. The first 3D classification step excluded particles in classes without RNAP density, while 73,254 particles from classes with some RNAP density were further classified into the 16 classes shown here. Two classes (blue) showed strong RNAP signal and are termed 'tightly-coupled' expressomes:

these particles were used for all subsequent analysis. Numerous classes contained weak RNAP signal (pink) and are termed ‘loosely-coupled’ expressomes. Further classification of these particles did not yield reconstructions with interpretable RNAP density. **(B)** Representative 2D class averages. **(C)** Improvement in cryo-EM maps by multi-body refinement. Initial reconstruction from tightly-coupled expressome particles (left, shown low-pass filtered to 8 Å) was used to generate masks for the ribosome and RNAP regions (blue and red respectively). The resulting maps of improved resolution are shown following rigid-body fitting into the consensus positions of the initial reconstruction. **(D)** Fourier shell correlation plots for gold standard refinements. Labels indicate the expressome region within the mask used for each refinement. **(E)** Local resolution map of the ribosome and RNAP regions within the NusG-coupled expressome. The map refined with a mask around the 70S ribosome ranges from ~3.4 Å in the core to ~4.5 Å for regions on the surface. The map refined with a mask around RNAP ranges from ~6 Å in the core to >12 Å for domains typically identified as mobile.

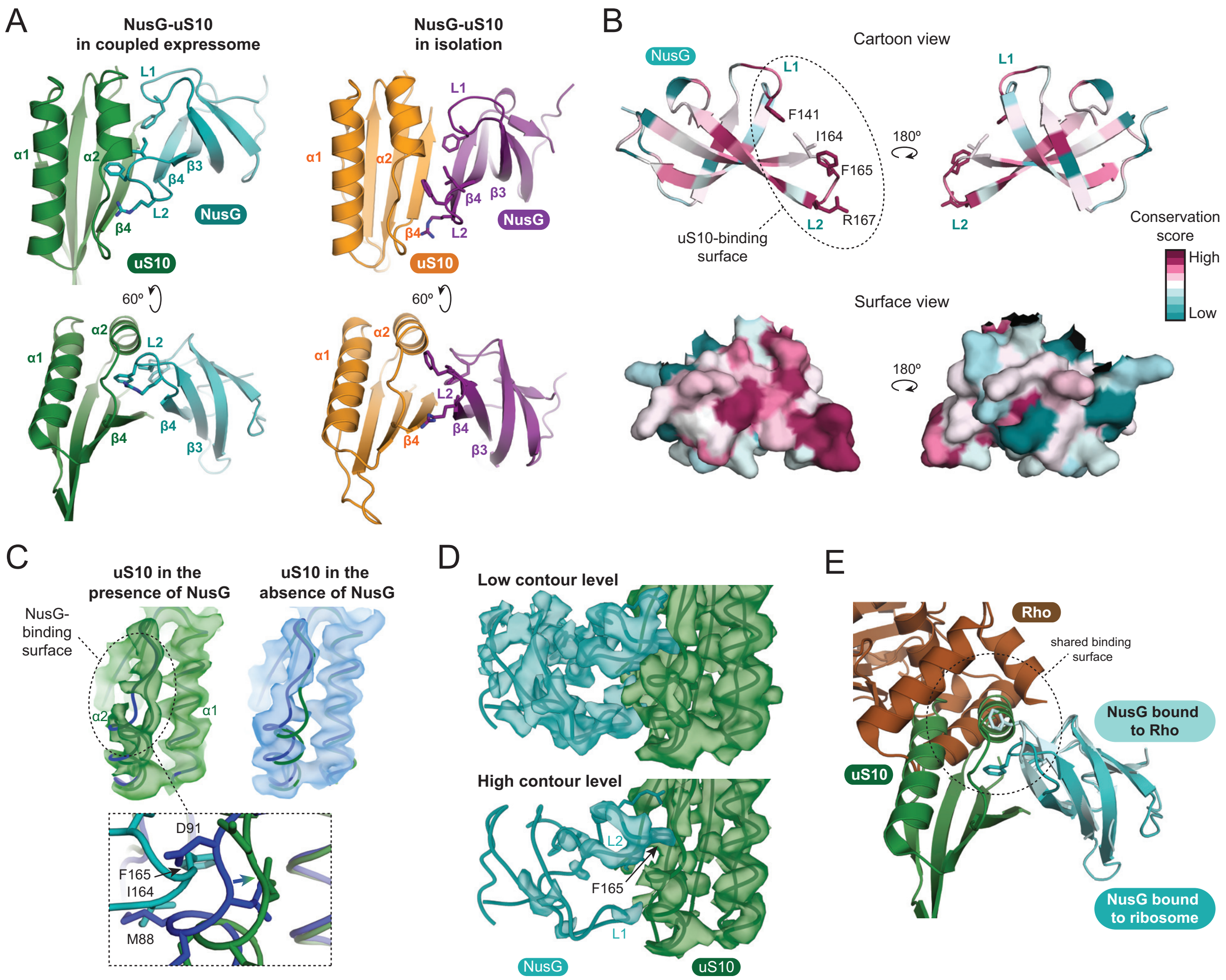


**Figure S5**

**Figure S5:** Details of the NusG-uS10 interaction in the NusG-coupled expressome. **(A)** The interaction between uS10 and NusG in the context of the NusG-coupled expressome (left) is similar but not identical to the interaction of isolated proteins (right; PDB code 2KVQ) (*7*). Rotation of the NusG-CTD relative to uS10 allows NusG loop L2 to insert into a hydrophobic pocket of uS10 formed by movement of helix α2 and strand β4. **(B)** Per-residue evolutionary conservation of *E. coli* NusG-CTD shows the uS10-binding region (dashed outline) is highly conserved. Calculations are based on alignment of 22 bacterial sequences selected from diverse phyla. **(C)** Electron density maps with superimposed backbone coordinates showing the different positions of uS10 helix α2 in NusG-coupled expressome (top left, green) and uncoupled expressome (top right, blue). This movement is required to accommodate NusG (bottom) and avoid clashes: NusG(F165)-uS10(D91) and NusG(I164)-uS10(M88). **(D)** NusG density is most well resolved for residues that contact uS10. While the entire NusG-CTD domain is visible at low map contour levels, only residues within loop L2 are visible at high contour levels. This indicates flexibility within the NusG-CTD domain. **(E)** Structural overlay of the NusG-uS10 complex within the NusG-coupled expressome with the NusG-Rho complex involved in transcription termination (*15*) showing the shared interface (dashed outline).

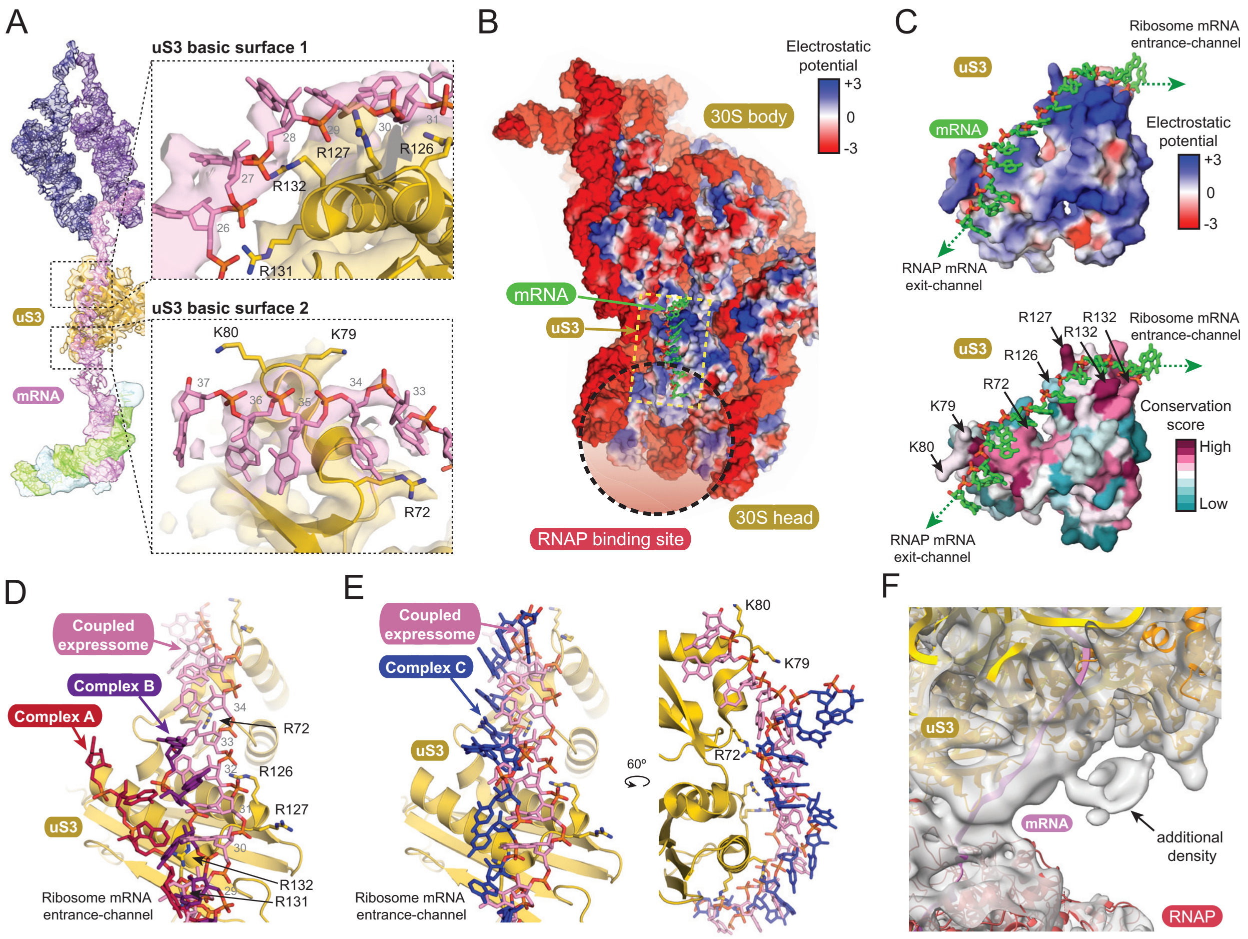


**Figure S6**

**Figure S6:** Details of the mRNA path in the NusG-coupled expressome. **(A)** The mRNA path from synthesis to decoding shown with Segmented cryo-EM density map (filtered to 4 Å resolution) and superimposed model for the NusG-coupled expressome. In the structure, mRNA residues 16-18 are within the ribosomal P-site and base-paired with tRNA$^{fMet}$, residues 19-21 are within the ribosomal A-site and base-paired with Phe-tRNA$^{Phe}$. mRNA residues 22-26 are within the ribosomal mRNA entry channel and contact the interface between ribosomal proteins uS3 and uS4. Inset 'uS3 basic surface 1' (top right) shows mRNA residues 27-31 on the surface of ribosomal protein uS3 at the edge of the mRNA entrance-channel positioned close to four arginines: R126, R127, R131 and R132. This basic surface likely coordinates the negatively charged phosphate backbone. Inset 'uS3 basic surface 2' (bottom right) shows mRNA residues 33-36 on the surface of uS3 closest to RNAP that contains three basic residues: R72, K79 and K80. These mRNA bases appear to lie within a hydrophobic groove. mRNA residues 30-32 are between these surfaces and, without contacts to the surface, are poorly ordered. mRNA residues 37-44 are within the RNAP exit-channel and poorly ordered. mRNA residues 45-53 are within RNAP and base-paired with template DNA. **(B)** Electrostatic surface potential of the 30S subunit of the *E. coli* ribosome indicating the location of positively charged surface of uS3 that connects the mRNA entrance-channel to the mRNA exit-channel of RNAP. **(C)** The surface of uS3 that contacts mRNA is highly conserved and positively charged. Electrostatic surface potential (left) and per-residue evolutionary conservation (right) are shown for *E. coli* ribosomal protein uS3. Conservation was calculated based on alignment of 29 bacterial sequences selected from diverse phyla. The superimposed mRNA path (green) shows the uS3 surfaces involved in contact. **(D)** Comparison of the mRNA path of the NusG-coupled expressome with that of mRNA-ribosome complexes stabilized by introduction of a 3' hairpin (*19*). Different mRNA paths were observed for the two molecules in the asymmetric unit and are named 'Complex A' and 'Complex B' in accordance with the previous publication. Of the basic residues outside the entrance-channel that likely contact mRNA in the NusG-coupled expressome, only R131 and R132 were also identified in previous structures. **(E)** Comparison of the mRNA path of the NusG-coupled expressome with that of a mRNA-ribosome complex stabilized with a frameshift-inducing mRNA hairpin ('Complex C') (*48*). The mRNA hairpin lies on the ribosome surface in approximately the same region as the mRNA path in the NusG-coupled expressome, likely due to electrostatic attraction (note the mRNA hairpin model shown has been truncated to the region closest to uS3 for clarity). In particular, uS3 residue 72 appears to make contact in both structures. **(F)** Cryo-EM map low-pass filtered to 8 Å shows additional density connected to the intervening mRNA at low map contour levels. The map quality is insufficient for confident assignment. The location is approximately consistent, however, with electron density previously assigned to ribosomal protein bS1 (*49*), supported by cross-linking mass spectrometry (*50*). The density does not directly connect to either the ribosome or RNAP, consistent with it arising from an RNA-binding domain such as a bS1 OB domain.

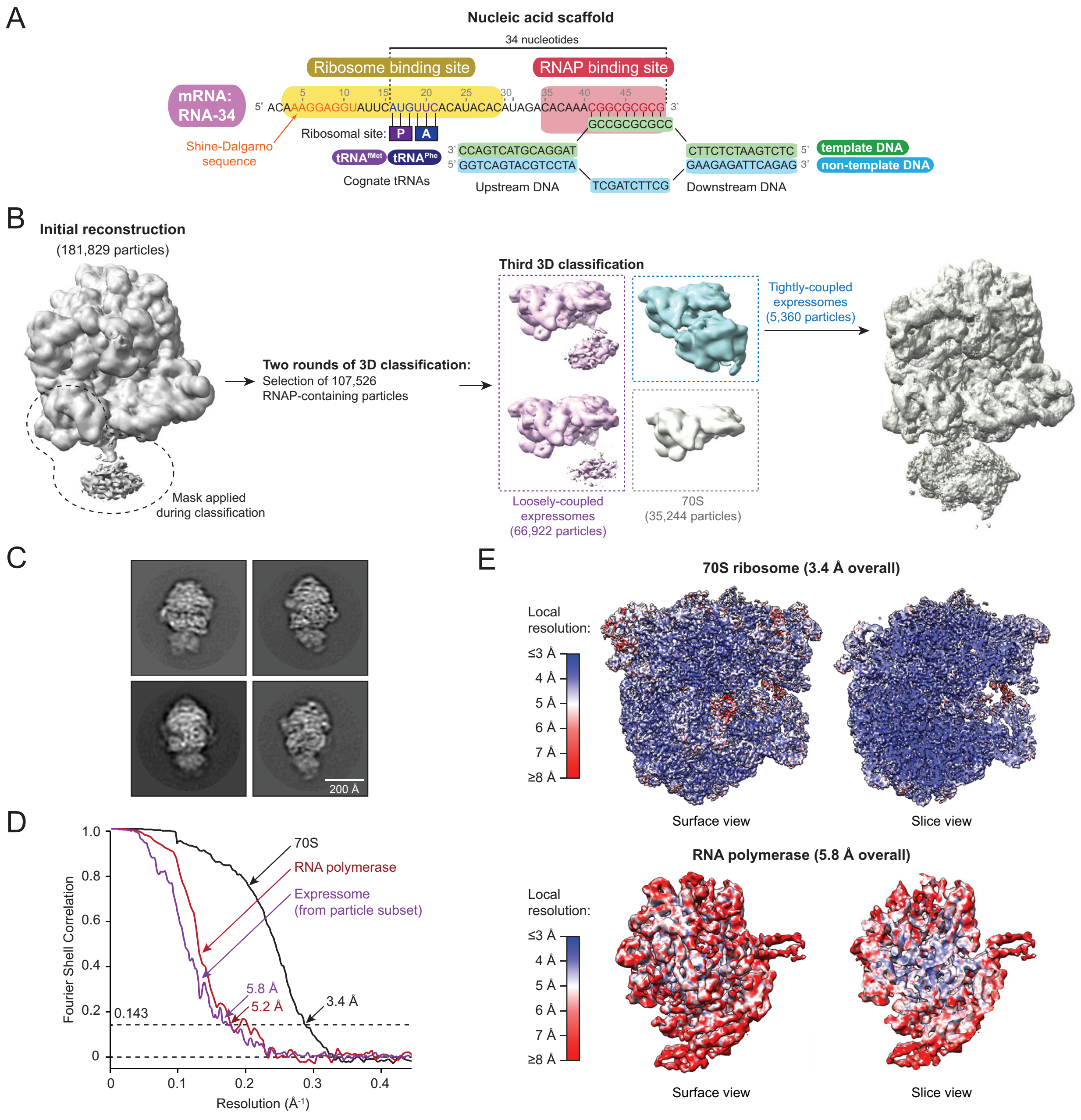


**Figure S7**

**Figure S7:** Assembly and cryo-EM of the collided expressome. **(A)** Schematic of nucleic acid scaffold used to assemble collided expressome complexes for cryo-EM. Key sequence features of the mRNA (RNA-34) are colored: Shine-Dalgarno sequence for ribosome binding (orange), codons for tRNA binding (blue and purple), sequence complementary to template DNA (red). mRNAs are named according to the number of nucleotides between, and including, the ribosomal P-site and the RNA-DNA duplex. **(B)** 3D classification scheme: Initial 2D and 3D classification produced 181,829 particles which were a mixture of expressomes and ribosomes. Expressome particles were selected by two methods. First, 45,774 particles were selected by signal subtraction of the ribosome as described in Fig S1E, and were used for high-resolution ribosome and RNAP reconstructions and quantification of relative orientation. Second, more stringent selection by 3D classification (shown here) yielded 5,360 particles that produced a well-defined expressome reconstruction. A significant number of particles are therefore in state different from the consensus presented in Fig 3, and the sample as a whole is best understood with consideration of the relative orientation data (Fig 4A and S8B). **(C)** Representative 2D class averages for the 5,360 particles. **(D)** Fourier shell correlation plots for gold standard refinements of the 45,774 particles. Labels indicate the expressome region within the mask used for each refinement. **(E)** Local resolution map of the ribosome and RNAP regions. The map refined with a mask around the 70S ribosome ranges from ~3.4 Å in the core to ~4.5 Å for regions on the surface. The map refined with a mask around RNAP ranges from ~6 Å in the core to >12 Å for domains typically identified as mobile.

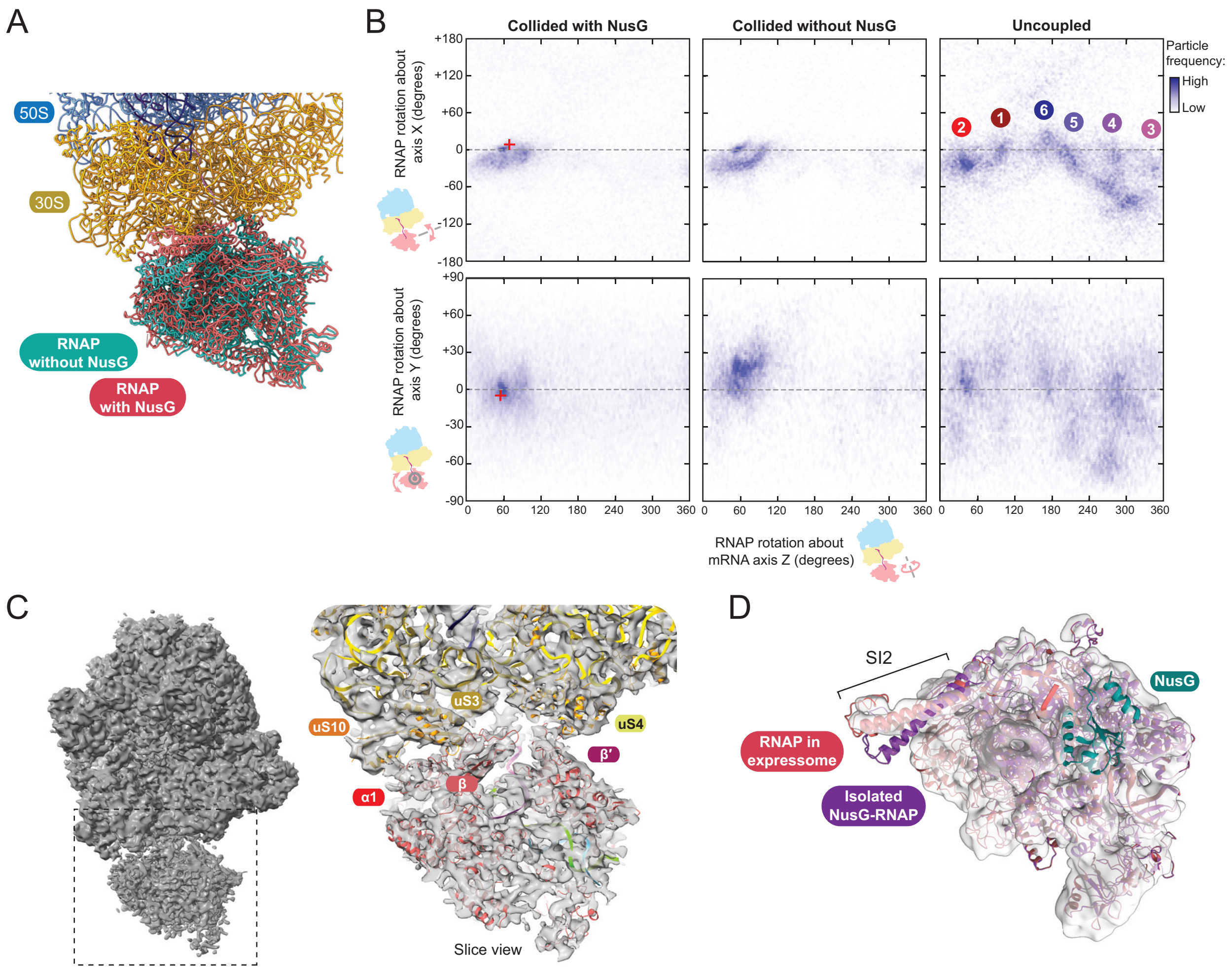


**Figure S8**

**Figure S8:** Features of the collided expressome. **(A)** Comparison of collided expressome structures in the presence (red) and absence (cyan) of NusG. **(B)** Heatmaps of the position of RNAP relative to the ribosome in the collided expressome with NusG (RNA-34) (left), collided expressome without NusG (RNA-34) (middle), and uncoupled expressome without NusG (RNA-38). Note that the spatial definition of the Euler angles representing relative orientation differ from Fig 1C and S3A in the direction of rotation about the mRNA axis and an offset of 60˚. The measured orientation of RNAP relative to the ribosome reported for the collided expressome is indicated as a red cross for comparison (*9*). The top left panel is reproduced from Figure 4A. **(C)** Cryo-EM map of the collided expressome without focused refinement (left), and slice view of the interaction interface of RNAP with the ribosome indicating map quality (right). **(D)** RNAP sequence insertion 2 (SI2) of subunit β adopts a different conformation in the expressome (red) compared with the isolated RNAP-NusG complex (purple; PDB code 6C6U) (*13*). Map of RNAP from focused refinement shown with transparency.

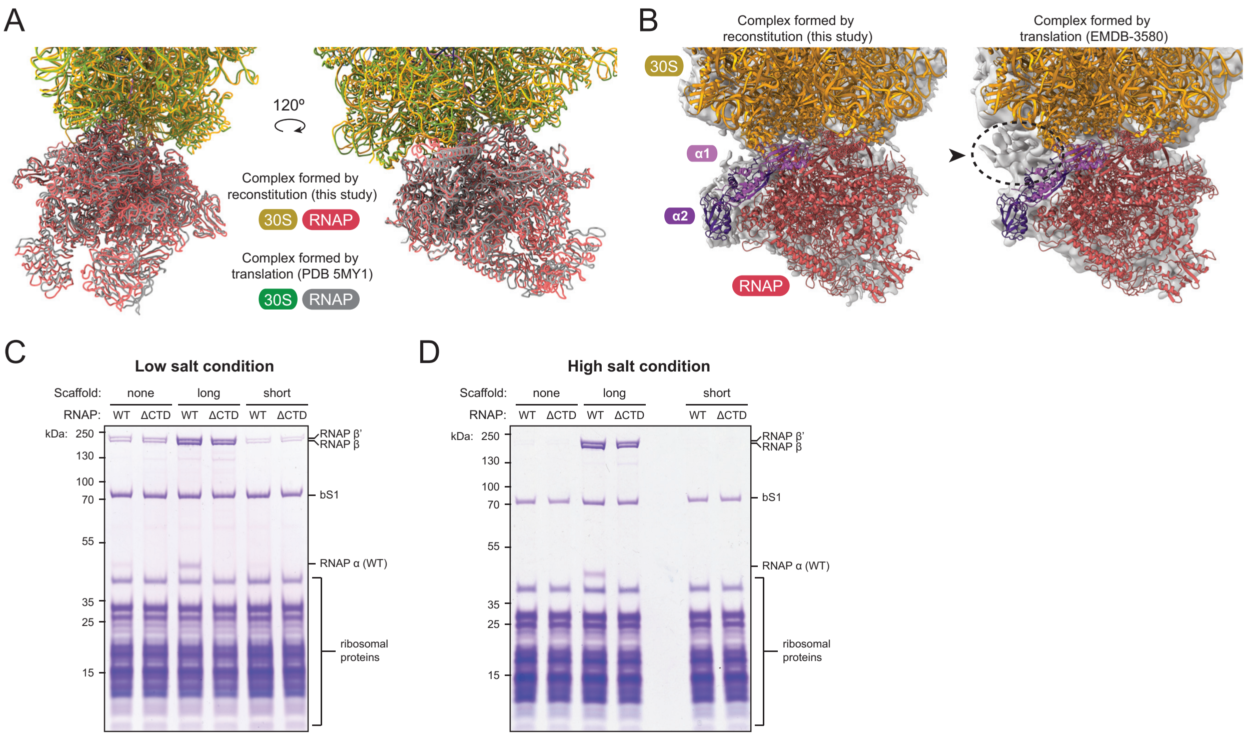


**Figure S9**

**Figure S9:** The collided expressome formed by reconstitution resembles that formed by translation. **(A)** Close structural resemblance between our expressome model (yellow and red) with a published model obtained by collision of ribosomes with stalled RNAP using an in vitro translation kit (green and grey) (*9*). Superposition is based on alignment of 16S rRNA. **(B)** Reconstructions in this study (left) did not indicate the additional density that was previously assigned to the C-terminal domains of the RNAP α-subunits (α-CTDs) (arrow, EMDB code 3580) (*9*). **(C-D)** An RNAP variant lacking the α-CTD domains ('ΔCTD') co-purifies with ribosomes similarly to wild-type RNAP ('WT'), consistent with no major role of the α-CTDs in expressomes formed by reconstitution. Components were mixed either with an mRNA long enough to allow ribosome binding ('long'), or not ('short'), or no nucleic acids ('none'). Assembly and gradient purification were performed in either (C) low salt (50 mM NaCl, 25 mM $MgCl_2$), or (D) high salt (120 mM KOAc, 20 mM $Mg(OAc)_2$, 10 mM $NH_4Cl$). Binding of RNAP to ribosomes is independent of ionic strength with the long mRNA, while complexes stable enough for gradient centrifugation in the presence of the short mRNA, or no scaffold were only observed in the lower salt condition.

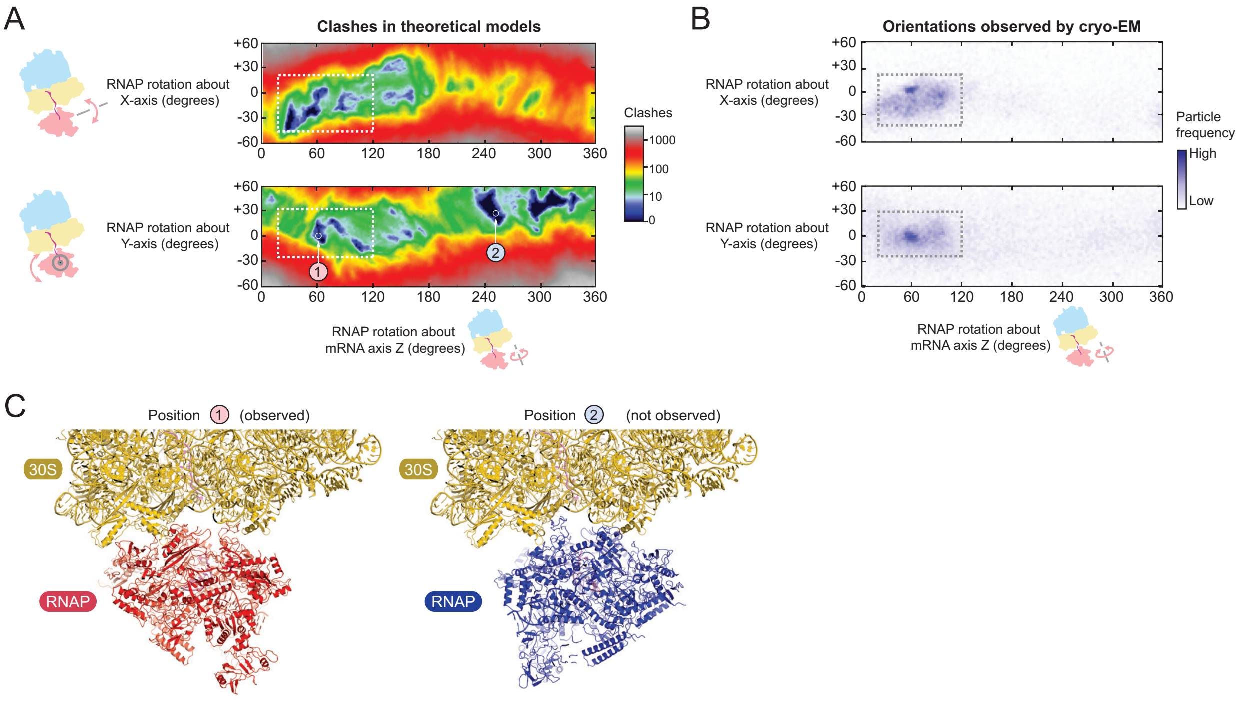


**Figure S10**

**Figure S10:** Collided expressome formation is driven by structural complementarity. **(A)** Heatmaps showing orientations of RNAP relative to the ribosome that avoid clashes. A series of atomic models were generated by varying the RNAP:ribosome relative orientation by incremental rotation of RNAP-NusG from its position in the collided expressome (defined here by rotational coordinates z=60,y=0,x=0). For each resulting model, clashes between backbone atoms of RNAP and the ribosome were counted. Blue regions indicate expressome configurations clashes do not occur due to structural complementarity. Two cross-sections of the three-dimensional rotational space are shown. **(B)** Heatmaps of the orientation of RNAP relative to the ribosome observed by cryo-EM, reproduced from Figure S8B for comparison with theoretical expressome models (dashed boxes). **(C)** Expressome models corresponding to points ‘1’ (left) and ‘2’ (right) indicated in panel A. While both models are sterically allowed, configurations similar to ‘1’ were observed by cryo-EM, while those similar to ‘2’ were not. Reduced contact area of RNAP with ribosome in ‘2’ may disfavor this alternative binding arrangement.

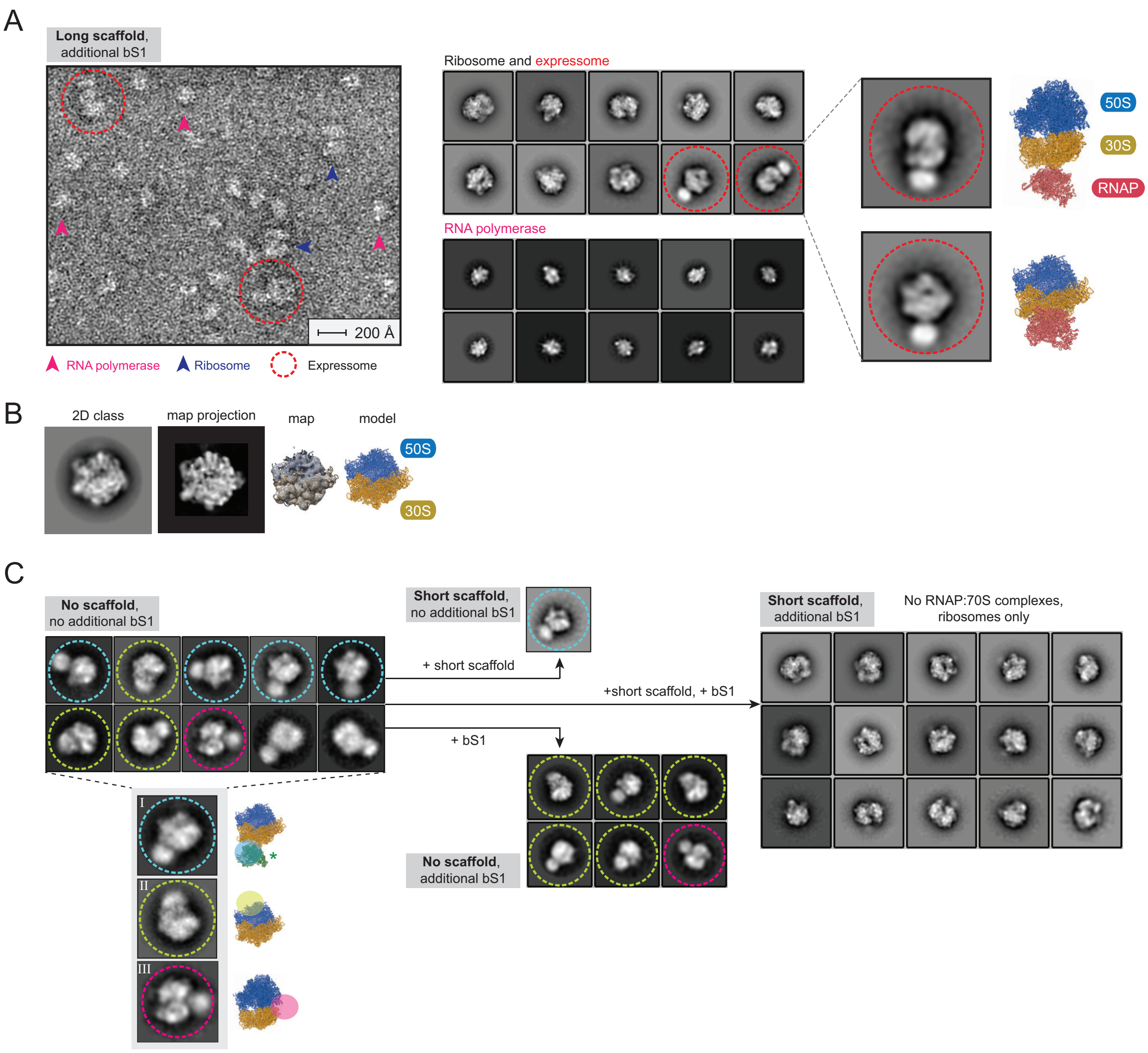


**Figure S11**

**Figure S11:** Analysis of 70S-RNAP complexes by negative Stain EM. **(A)** Expressomes formed by assembly on a long scaffold that supports ribosome binding (RNA-34) imaged by negative stain EM. Representative micrograph (left), 2D class averages grouped into those that contain ribosomes (center top) and those that contain RNAP (center bottom), and selected expressome classes (right) with the corresponding view of the collided expressome model obtained by cryo-EM. **(B)** The direction of view in the 2D classes presented in Fig 4D (reproduced here) was determined by similarity to a projection of the negative stain 3D reconstruction shown here. The assignment is also consistent with the expected position of RNAP in the expressome. **(C)** Core

RNAP-70S complex 2D class averages (left). Classes with the view corresponding to panel B presented in Fig 4D. Saturation of ribosomes by addition of ribosomal protein bS1 abolished occupancy of site I (a site consistent in position with the published core RNAP-30S complex structure (PDB code 6AWB) (*10*)), and only complexes with occupied site II could be observed (top center). Addition of a nucleic acid scaffold with short mRNA abolished site II without affecting site I (bottom center). Addition of both, bS1 and a nucleic acid scaffold with short mRNA abolished occupancy of both alternative sites and no expressomes formed either (right). The dependence on mRNA for complex formation is supported by comparison of short scaffold with bS1 (right) with long scaffold with bS1 (panel A). Attempts to reconstruct RNAP-ribosome complexes in 3D from negative stain data were unsuccessful. This is likely due to limited number of particles, their preferred orientation on grids, and heterogeneity in the 70S-RNAP interfaces.

**Table S1. Refinement and model statistics for four reconstructions of supramolecular assemblies of *E. coli* RNAP and the 70S ribosome**

| Data collection | Uncoupled Expressome | | Coupled Expressome (excess NusG) | | Collided Expressome (excess NusG) | | Collided Expressome ( no NusG) | |
|---|---|---|---|---|---|---|---|---|
| Particles | 32195 | | 15327 | | 45774 | | 18552 | |
| Pixel size (Å) | 1.052 | | 1.052 | | 1.075 | | 1.075 | |
| Defocus range (um) | 0.7-3.5 | | 0.7-3.5 | | 0.7-3.5 | | 0.7-3.5 | |
| Voltage (kV) | 300 | | 300 | | 300 | | 300 | |
| Electron dose ($e^{-}$ Å$^{-2}$) | 42 | | 42 | | 50 | | 51 | |
| **Model composition** | 70S | RNAP | 70S | RNAP | 70S | RNAP | 70S | RNAP |
| Non-hydrogen atoms | 146829 | 25527 | 150187 | 26949 | 145833 | 26949 | 146319 | 26159 |
| Protein residues | 5692 | 3101 | 6110 | 3273 | 5642 | 3273 | 5628 | 3175 |
| RNA bases | 4739 | 11 | 4749 | 14 | 4729 | 14 | 4739 | 14 |
| DNA bases | - | 51 | - | 61 | - | 61 | - | 61 |
| Ligands ($Zn^{2+}$/$Mg^{2+}$) | 2/539 | 2/1 | 2/539 | 2/1 | 1/197 | 2/1 | 1/533 | 2/1 |
| **Focused Refinement** | 70S | RNAP | 70S | RNAP | 70S | RNAP | 70S | RNAP |
| Nominal resolution (Å) | 3.0 | 3.8 | 3.4 | 7.6 | 3.4 | 5.8 | 3.3 | 5.0 |
| Map sharpening B-factor (Å$^2$) | -58.3 | -133.0 | -70.2 | -136.5 | -75.7 | -175.2 | -100.9 | -198.6 |
| Map cross-correlation (within mask) | 0.88 | 0.83 | 0.83 | 0.75 | 0.85 | 0.75 | 0.86 | 0.67 |
| Average B factor protein (Å$^2$) | 37.4 | 75.1 | 102.8 | 431.6 | 85.3 | 235.8 | 67.5 | 196.2 |
| Average B factor nucleotide (Å$^2$) | 46.6 | 140.8 | 104.8 | 440.0 | 91.4 | 256.5 | 78.1 | 189.6 |
| **RMS deviations** | | | | | | | | |
| Bond lengths (Å) | 0.01 | 0.01 | 0.01 | 0.01 | 0.01 | 0.01 | 0.01 | 0.01 |
| Bond angles (°) | 0.777 | 0.949 | 0.756 | 1.043 | 0.826 | 1.042 | 0.79 | 1.043 |
| **Ramachandran plot** | | | | | | | | |
| Favored (%) | 93.54 | 82.85 | 90.37 | 91.66 | 88.71 | 91.69 | 91.67 | 91.63 |
| Allowed (%) | 6.36 | 17.12 | 9.48 | 8.22 | 11.18 | 8.19 | 8.19 | 8.24 |
| Outliers (%) | 0.11 | 0.03 | 0.15 | 0.12 | 0.11 | 0.12 | 0.14 | 0.13 |
| **Validation** | | | | | | | | |
| Molprobity Score | 2.44 | 3.38 | 2.97 | 1.87 | 3.15 | 1.89 | 2.88 | 1.87 |
| Molprobity Clash score | 7.93 | 22.87 | 12.46 | 6.66 | 16.39 | 7.04 | 13.27 | 6.71 |
| Rotamer outliers (%) | 5.91 | 12.46 | 11.97 | 0.29 | 13.02 | 0.3 | 9.63 | 0.3 |
| **Validation (RNA/DNA)** | | | | | | | | |
| Correct sugar puckers (%) | 99.64 | 100 | 99.62 | 100 | 99.56 | 100 | 99.68 | 100 |
| Good backbone conformation (%) | 84.2 | 54.55 | 81.83 | 64.29 | 80.78 | 64.29 | 82.01 | 64.29 |

**Table S1:** Summary of data collection and refinement statistics. Ribosome models were real-space refined with Phenix (*36*) (rigid body refinement, global minimization, and ADP refinement) against focused reconstructions for all four structures. RNAP was refined in the uncoupled expressome was real-space refined (rigid body refinement, global minimization, and ADP refinement) or docked as a rigid body using a published model (6C6U) (*13*) and refined as a rigid body in Phenix (rigid body refinement, and ADP refinement).

**Movie S1:** Movement of RNA polymerase (RNAP) relative to the 70S ribosome in the uncoupled expressome. The orientations of RNAP relative to the ribosome was quantified (bottom right), and expressome models generated from particles in each of six clusters. Two views of the expressome (top) show the continuous movement of RNAP relative to the ribosome calculated by interpolation (RNAP red, 30S yellow, 50S blue). A magnified view of the 30S surface that contacts RNAP is shown coloured by height (bottom left). The zinc finger of the β′ subunit (β′-ZF, purple) is the closest region of RNAP to the ribosome surface in all states. The β′-ZF moves within a depression from the mRNA entrance-channel to a site bounded by ribosomal proteins uS3 and uS10 on the 30S head domain. One state is consistent with the formation of a bridge by the two domains of transcription factor NusG (C-terminal domain 'CTD' blue, N-terminal domain 'NTD' cyan).

**Movie S2:** Movement of RNA polymerase (RNAP) relative to the 70S ribosome in the NusG-coupled expressome. The orientations of RNAP relative to the ribosome was determined by cryo-EM multi-body refinement. Two views of the expressome (top) show the continuous movement of RNAP relative to the ribosome calculated by interpolation of models derived from two principal components (RNAP red, 30S yellow, 50S blue). A magnified view of the 30S surface that contacts RNAP is shown coloured by height (bottom left). The zinc finger of the β′ subunit (β′-ZF, purple) is the closest region of RNAP to the ribosome surface. It is contained on three sides by ribosomal proteins uS3, uS10 and helix 33 of 16S ('h33'). The NusG C-terminal domain 'CTD' is bound to uS10, while the N-terminal domain 'NTD' is bound to RNAP. The eight amino acid linker connecting these domains is poorly ordered, and is shown here as a dashed line joining residues Q117 and T126. Limited extension of the NusG linker likely contributes to movement of RNAP.

**Movie S3:** Movement of RNA polymerase (RNAP) relative to the 70S ribosome in the collided expressome. The orientations of RNAP relative to the ribosome was quantified (bottom right), and models were generated that correspond to the approximate limits of observed positions (±45° about horizontal axis, ±15° about vertical axis). Two views of the expressome (top) show the continuous movement of RNAP relative to the ribosome calculated by interpolation (RNAP red, 30S yellow, 50S blue). A magnified view of the 30S surface that contacts RNAP is shown coloured by height (bottom left). The 'footprint' of RNAP, corresponding to regions close to the ribosome surface, are shown in cartoon representation (purple). The point of mRNA exit from RNAP is indicated by a pink sphere. Complementarity between the convex RNAP surface and the concave ribosome surface constrain RNAP movement.

### References


1. R. Byrne, J. G. Levin, H. A. Bladen, M. W. Nirenberg, The *in vitro* formation of a DNA-ribosome complex. *Proc Natl Acad Sci USA*. **52**, 140–148 (1964).
2. O. L. Miller, B. A. Hamkalo, C. A. Thomas, Visualization of bacterial genes in action. *Science*. **169**, 392–395 (1970).
3. C. Yanofsky, Attenuation in the control of expression of bacterial operons. *Nature*. **289**, 751–758 (1981).
4. J. P. Richardson, Preventing the synthesis of unused transcripts by Rho factor. *Cell*. **64**, 1047–1049 (1991).
5. S. Proshkin, A. R. Rahmouni, A. Mironov, E. Nudler, Cooperation between translating ribosomes and RNA polymerase in transcription elongation. *Science*. **328**, 504–508 (2010).
6. M. Zhu, M. Mori, T. Hwa, X. Dai, Disruption of transcription-translation coordination in Escherichia coli leads to premature transcriptional termination. *Nat Microbiol*. **54**, 1–10 (2019).
7. B. M. Burmann *et al.*, A NusE:NusG Complex Links Transcription and Translation. *Science*. **328**, 501–504 (2010).
8. S. Saxena *et al.*, Escherichia coli transcription factor NusG binds to 70S ribosomes. *Mol Microbiol*. **108**, 495–504 (2018).
9. R. Kohler, R. A. Mooney, D. J. Mills, R. Landick, P. Cramer, Architecture of a transcribing-translating expressome. *Science*. **356**, 194–197 (2017).
10. G. Demo *et al.*, Structure of RNA polymerase bound to ribosomal 30S subunit. *Elife*. **6**, 94 (2017).
11. H. Fan *et al.*, Transcription-translation coupling: direct interactions of RNA polymerase with ribosomes and ribosomal subunits. *Nucleic Acids Res.* **45**, 11043–11055 (2017).

12. D. Castro-Roa, N. Zenkin, In vitro experimental system for analysis of transcription-translation coupling. *Nucleic Acids Res*. **40**, e45–e45 (2012).

13. J. Y. Kang *et al.*, Structural Basis for Transcript Elongation Control by NusG Family Universal Regulators. *Cell*. **173**, 1650–1662.e14 (2018).

14. S. L. Sullivan, M. E. Gottesman, Requirement for E. coli NusG protein in factor-dependent transcription termination. *Cell*. **68**, 989–994 (1992).

15. M. R. Lawson *et al.*, Mechanism for the Regulated Control of Bacterial Transcription Termination by a Universal Adaptor Protein. *Mol Cell*. **71**, 911–922.e4 (2018).

16. M. Turtola, G. A. Belogurov, NusG inhibits RNA polymerase backtracking by stabilizing the minimal transcription bubble. *Elife*. **5** (2016), doi:10.7554/eLife.18096.

17. G. Vauquelin, S. J. Charlton, Exploring avidity: understanding the potential gains in functional affinity and target residence time of bivalent and heterobivalent ligands. *Br. J. Pharmacol.* **168**, 1771–1785 (2013).

18. S. Takyar, R. P. Hickerson, H. F. Noller, mRNA helicase activity of the ribosome. *Cell*. **120**, 49–58 (2005).

19. H. Amiri, H. F. Noller, Structural evidence for product stabilization by the ribosomal mRNA helicase. *RNA*. **25**, 364–375 (2019).

20. X. Qu *et al.*, The ribosome uses two active mechanisms to unwind messenger RNA during translation. *Nature*. **475**, 118–121 (2011).

21. C. L. Chan, R. Landick, The Salmonella typhimurium his operon leader region contains an RNA hairpin-dependent transcription pause site. Mechanistic implications of the effect on pausing of altered RNA hairpins. *J Biol Chem*. **264**, 20796–20804 (1989).

22. I. Gusarov, E. Nudler, The mechanism of intrinsic transcription termination. *Mol Cell*. **3**, 495–504 (1999).

23. T. Nakane, D. Kimanius, E. Lindahl, S. H. Scheres, Characterisation of molecular motions in cryo-EM single-particle data by multi-body refinement in RELION. *Elife*. **7**, 1485 (2018).

24. K. R. Andersen, N. C. Leksa, T. U. Schwartz, Optimized E. coli expression strain LOBSTR eliminates common contaminants from His-tag purification. *Proteins*. **81**, 1857–1861 (2013).

25. K.-A. F. Twist *et al.*, A novel method for the production of in vivo-assembled, recombinant Escherichia coli RNA polymerase lacking the α C-terminal domain. *Protein Sci*. **20**, 986–995 (2011).

26. X. Guo *et al.*, Structural Basis for NusA Stabilized Transcriptional Pausing. *Mol Cell*. **69**, 816–827.e4 (2018).

27. M. N. Vassylyeva *et al.*, Purification, crystallization and initial crystallographic analysis of RNA polymerase holoenzyme from Thermus thermophilus. *Acta Crystallogr D Biol Crystallogr*. **58**, 1497–1500 (2002).

28. D. Moazed, H. F. Noller, Transfer RNA shields specific nucleotides in 16S ribosomal RNA from attack by chemical probes. *Cell*. **47**, 985–994 (1986).

29. G. Blaha *et al.*, Preparation of functional ribosomal complexes and effect of buffer conditions on tRNA positions observed by cryoelectron microscopy. *Meth Enzymol*. **317**, 292–309 (2000).

30. A. R. Subramanian, Structure and functions of ribosomal protein S1. *Prog. Nucleic Acid Res. Mol. Biol.* **28**, 101–142 (1983).

31. R. Jünemann *et al.*, In vivo deuteration of transfer RNAs: overexpression and large-scale purification of deuterated specific tRNAs. *Nucleic Acids Res*. **24**, 907–913 (1996).

32. E. Cayama *et al.*, New chromatographic and biochemical strategies for quick preparative isolation of tRNA. *Nucleic Acids Res*. **28**, E64 (2000).

33. S. Q. Zheng *et al.*, MotionCor2: anisotropic correction of beam-induced motion for improved cryo-electron microscopy. *Nat Methods*. **14**, 331–332 (2017).

34. K. Zhang, Gctf: Real-time CTF determination and correction. *J. Struct. Biol.* **193**, 1–12 (2016).

35. J. Zivanov *et al.*, New tools for automated high-resolution cryo-EM structure determination in RELION-3. *Elife*. **7** (2018), doi:10.7554/eLife.42166.

36. D. Liebschner *et al.*, Macromolecular structure determination using X-rays, neutrons and electrons: recent developments in Phenix. *Acta Crystallogr D Struct Biol*. **75**, 861–877 (2019).

37. J. Noeske *et al.*, High-resolution structure of the Escherichia coli ribosome. *Nat Struct Mol Biol*. **22**, 336–341 (2015).

38. J. Y. Kang *et al.*, Structural basis of transcription arrest by coliphage HK022 nun in an Escherichia coli RNA polymerase elongation complex. *Elife*. **6** (2017), doi:10.7554/eLife.25478.

39. E. Schmitt, M. Panvert, S. Blanquet, Y. Mechulam, Crystal structure of methionyl-tRNAfMet transformylase complexed with the initiator formyl-methionyl-tRNAfMet. *EMBO J*. **17**, 6819–6826 (1998).

40. R. T. Byrne, A. L. Konevega, M. V. Rodnina, A. A. Antson, The crystal structure of unmodified tRNAPhe from Escherichia coli. *Nucleic Acids Res*. **38**, 4154–4162 (2010).

41. E. F. Pettersen *et al.*, UCSF Chimera--a visualization system for exploratory research and analysis. *J Comput Chem*. **25**, 1605–1612 (2004).

42. M. Selmer *et al.*, Structure of the 70S ribosome complexed with mRNA and tRNA. *Science*. **313**, 1935–1942 (2006).

43. R. M. Voorhees, A. Weixlbaumer, D. Loakes, A. C. Kelley, V. Ramakrishnan, Insights into substrate stabilization from snapshots of the peptidyl transferase center of the intact 70S ribosome. *Nat Struct Mol Biol*. **16**, 528–533 (2009).

44. N. Fischer *et al.*, Structure of the E. coli ribosome-EF-Tu complex at <3 Å resolution by Cs-corrected cryo-EM. *Nature*. **520**, 567–570 (2015).

45. P. Emsley, K. Cowtan, Coot: model-building tools for molecular graphics. *Acta Crystallogr D Biol Crystallogr*. **60**, 2126–2132 (2004).

46. N. R. James, A. Brown, Y. Gordiyenko, V. Ramakrishnan, Translational termination without a stop codon. *Science*. **354**, 1437–1440 (2016).

47. J. B. Heymann, M. Chagoyen, D. M. Belnap, Common conventions for interchange and archiving of three-dimensional electron microscopy information in structural biology. *J. Struct. Biol.* **151**, 196–207 (2005).

48. Y. Zhang, S. Hong, A. Ruangprasert, G. Skiniotis, C. M. Dunham, Alternative Mode of E-Site tRNA Binding in the Presence of a Downstream mRNA Stem Loop at the Entrance Channel. *Structure*. **26**, 437–445.e3 (2018).

49. A. B. Loveland, A. A. Korostelev, Structural dynamics of protein S1 on the 70S ribosome visualized by ensemble cryo-EM. *Methods*. **137**, 55–66 (2018).

50. M. A. Lauber, J. Rappsilber, J. P. Reilly, Dynamics of ribosomal protein S1 on a bacterial ribosome with cross-linking and mass spectrometry. *Mol. Cell Proteomics*. **11**, 1965–1976 (2012).
